# *Toxoplasma* ceramide synthases: a curious case of gene duplication, divergence and key functionality

**DOI:** 10.1101/2022.01.05.475179

**Authors:** Zisis Koutsogiannis, John G. Mina, Christin A. Albus, Matthijs A. Kol, Joost C. M. Holthuis, Ehmke Pohl, Paul W. Denny

## Abstract

*Toxoplasma gondii* is an obligate, intracellular eukaryotic apicomplexan protozoan parasite that can cause foetal damage and abortion in both animals and humans. Sphingolipids have indispensable functions as signaling molecules and are essential and ubiquitous components of eukaryotic membranes that are both synthesized and scavenged by the Apicomplexa. Ceramide is the precursor for all sphingolipids, and here we report the identification, localisation and analyses of the *Toxoplasma* ceramide synthases *Tg*CerS1 and *Tg*CerS2 and, using a conditional gene regulation approach, establish their roles in pathogenicity and parasite fitness. Interestingly, we observed that whilst *Tg*CerS1 was a fully functional orthologue of the yeast Lag1p capable of catalysing the conversion of sphinganine to ceramide, in contrast *Tg*CerS2 was catalytically inactive. Furthermore, genomic deletion of *Tg*CerS1 using CRISPR/Cas-9 led to viable but slow growing parasites indicating its importance but not indispensability. In contrast, genomic knock out of *Tg*CerS2 was only accessible utilising the rapamycin-inducible Cre recombinase system. Surprisingly, the results demonstrated that this ‘pseudo’ ceramide synthase, *Tg*CerS2, has an even greater role in parasite fitness than its catalytically active orthologue (*Tg*CerS1). Phylogenetic analyses indicated that, as in humans and plants, the ceramide synthase isoforms found in *Toxoplasma* and other Apicomplexa arose through gene duplication. However, in the Apicomplexa the duplicated copy subsequently evolved into a non-functional ‘pseudo’ ceramide synthase. This arrangement is unique to the Apicomplexa and further illustrates the unusual biology that characterize these protozoan parasites, a feature that could potentially be exploited in the development of new antiprotozoals.

**Author Summary:** Sphingolipids, essential and ubiquitous lipids in the Eukaryota, are both synthesized and scavenged by the parasitic apicomplexan protozoa, including *Toxoplasma gondii*. Ceramide is the precursor for all sphingolipids and here we report the identification, localisation and analyses of the *Toxoplasma* ceramide synthases *Tg*CerS1 and *Tg*CerS2. Surprisingly, whilst *Tg*CerS1 was fully functional, catalysing the conversion of sphinganine to ceramide, *Tg*CerS2 was catalytically inactive. However, we demonstrated that this ‘pseudo’ ceramide synthase has an even greater role in parasite fitness than the catalytically active *Tg*CerS1. Phylogenetic analyses indicated that these isoforms arose through gene duplication and the duplicated copy subsequently evolved into the ‘pseudo’ ceramide synthase. This arrangement is unique to the Apicomplexa and further illustrates the highly unusual biology that characterizes these protozoan parasites, a feature that could potentially be exploited for the development of new antiprotozoals.

## Introduction

The Apicomplexa are one of the four groups of infectious protozoa [1], and comprise a variety of pathogens of both animals and humans: *Cryptosporidium spp.* (diarrhoea); *Eimeria spp.* (coccidiosis in poultry and cattle); *Theileria spp.* (East Coast Fever in cattle); *Plasmodium spp.,* including *P. falciparum* the causative agent of severe malaria [2]; and *Toxoplasma gondii*, leading to toxoplasmosis. *Toxoplasma* is an obligate intracellular parasite with the ability to invade and colonise a broad range of nucleated vertebrate cells, therefore toxoplasmosis is one of the most prevalent infections and is estimated to affect 2-3 billion people worldwide [3]. The parasite has a highly complex lifecycle, Felidae being the definitive host and both proliferative (tachyzoite) and encysted (bradyzoite) forms in humans and other animals [4]. While the tachyzoite form can cause symptomatic disease, the bradyzoite stage is usually asymptomatic although, encysted in the brain, it has been proposed to be associated with numerous psychiatric disorders including schizoprenia [5]. Additionally, the presence of the dormant, encysted bradyzoites represents a continuous threat in immunocompromised individuals, particularly HIV patients, those receiving anti-cancer chemotherapy and organ transplant recipients [6], as *Toxoplasma* can transform into proliferative tachyzoite forms which cause symptomatic diseases such as Toxoplasmic Encephalitis (TE). Furthermore, *Toxoplasma* infection is a significant cause of congenital defects and subsequent abortions in both economically important domestic animals [4] and humans [6].

Treatment regimens for toxoplasmosis patients are limited and have remained largely unchanged since the 1950s [7]. Furthermore, there is no treatment available for chronic infection with encysted bradyzoite forms [8]. To this end efforts are focused on identifying new potential drug targets. Recent studies have shown major differences in the lipid, particularly sphingolipid, profiles of apicomplexans compared to the host [3, 9, 10]. Previously, sphingolipid biosynthesis has been validated as a drug target against the kinetoplastid protozoan parasites *Leishmania* [11–13] and *Trypanosoma* [14], as well as fungal pathogens [15–17]. Sphingolipids are ubiquitous amphipathic plasma membrane lipids involved in a myriad of signalling processes in all eukaryotes, including *Toxoplasma* [18]. However, the parasite harbours sphingolipid species that are absent in the mammalian host, including the phosphosphingolipid inositol phosphorylceramide [9]. Additionally, whilst *Toxoplasma* have been shown to possess sphingomyelin (the primary mammalian phosphosphingolipid), they have also been demonstrated to have a relatively high level of ethanolamine phosphorylceramide, which is only found as a trace phosphosphingolipid in the host cell [3].

As an intracellular parasite, *Toxoplasma* resides within a specialized parasitophorous vacuole (PV) formed immediately after invasion and delineated by the PV membrane [19]. However, *Toxoplasma* maintains a dynamic relationship with its host cell and small, soluble molecules such as purine [20] and aromatic amino acids [21] are able to cross the PV membrane by diffusion [22]. Additionally, *Toxoplasma* can efficiently salvage a range of key lipid species [23, 24]. For example, low-density lipoprotein (LDL)-derived cholesterol, for which the parasite is auxotrophic, is rapidly subverted to the PV via host microtubules [25, 26]. Some of the acquired lipids are then further metabolised by the parasite [27, 28]. Sphingolipids can also be scavenged from the host, [29–31], although host sphingolipid biosynthesis is non-essential for *Toxoplasma* proliferation [30, 31]. Indeed, the parasite retains a fully functional biosynthetic pathway [9, 29, 32] which is active throughout its lifecycle [33]. This indicated that *de novo* synthesis is important for parasitism and a potential target for therapeutic intervention [23].

Ceramide, the product of ceramide synthase (CerS or Lag1p in yeast), is the central molecule amongst the sphingolipid precursors which functions in a plethora of biological processes, including apoptosis, growth arrest and stress responses [34]. However, the first, rate-limiting stage in sphingolipid biosynthesis is mediated by serine palmitoyltransferase (SPT), which catalyses the condensation of L-serine and, typically, palmitoyl-CoA to form 3-keto-di-hydro-sphingosine (KDS or 3-keto-sphinganine) [35] in the endoplasmic reticulum (ER). Subsequently KDS is reduced and then *N*-acylated by CerS to form ceramide, the central metabolic hub of sphingolipid biosynthesis. Ceramide is then transported to the Golgi apparatus where it is converted into glycosphingolipids (GSLs) and, in mammals, sphingomyelin (SM) [36, 37].

Although *Toxoplasma* and the other apicomplexans maintain this pathway in principle, it is in many ways divergent when compared to the host cell machinery. For example, the first functionally characterised enzyme in the apicomplexan biosynthetic pathway was a functional orthologue of the yeast inositol phosphorylceramide (IPC) synthase, an enzyme with no mammalian equivalent [9, 38]. Furthermore, the apicomplexan SPT is, uniquely in the Eukaryota, of bacterial origin and presumed to have been acquired via lateral gene transfer [32]. However, the formal identification and characterisation of the apicomplexan CerS, the second key step in eukaryotic sphingolipid biosynthesis (Figure 1), was absent. Here we describe and characterise the *Toxoplasma Tg*CerS1 and *Tg*CerS2, which are representative of orthologues in the wider Apicomplexa. Utilising a combination of biochemical, genetic and bioinformatic tools we demonstrate that whilst *Tg*CerS1 is a functional CerS, *Tg*CerS2 lacks this activity *in vitro.* However, both are important for parasite fitness, with *Tg*CerS2 indicated to have a particularly important role. Phylogenetic analyses indicated that the apicomplexan ‘pseudo’ ceramide synthase, *Tg*CerS2, is the result of an ancient gene duplication event and a subsequent divergence that is unique to the Apicomplexa.

**Fig. 1.**
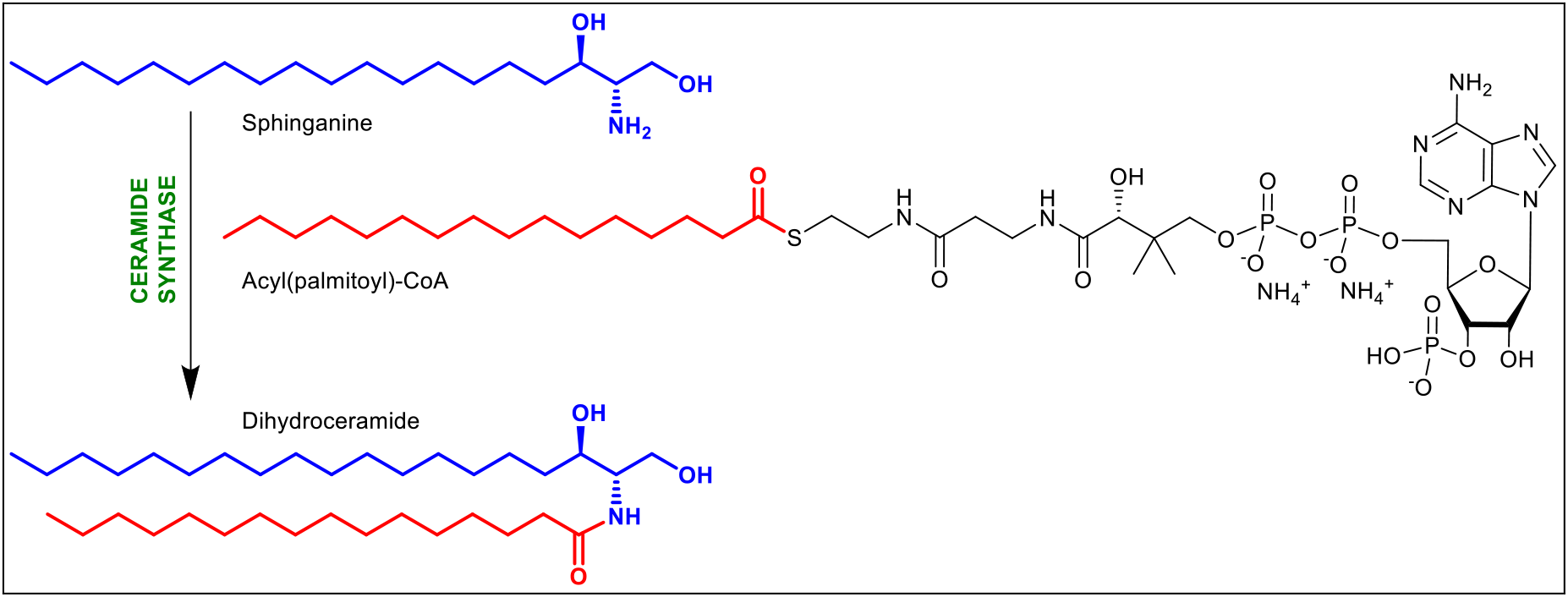
Substrates and products of ceramide synthase

## Results

### Identification of *Toxoplasma gondii* ceramide synthases

In all eukaryotic systems studied to date, ceramide synthases (CerS) are ER-resident integral membrane proteins that *N*-acylate di-hydrosphingosine (sphinganine) to produce di-hydroceramide which is then desaturated to form ceramide, the simplest sphingolipid and a key secondary messenger in numerous cellular pathways [39]. First identified as encoded by the longevity-assurance genes Lag1p and Lac1p in yeast [40], orthologues were subsequently found to be ubiquitous in the Eukaryota. All eukaryotes studied to date have been found to encode at least two CerS orthologues [41], with humans expressing six isoforms with each generating ceramides with a defined acyl chain length [41, 42]. All orthologues harbour the conserved 52 amino acid lag1p motif. Although the precise functionality of this domain remains unclear, 9 of the 11 canonical residues are essential for CerS activity (Figure 2B) [43].

**Fig. 2.**
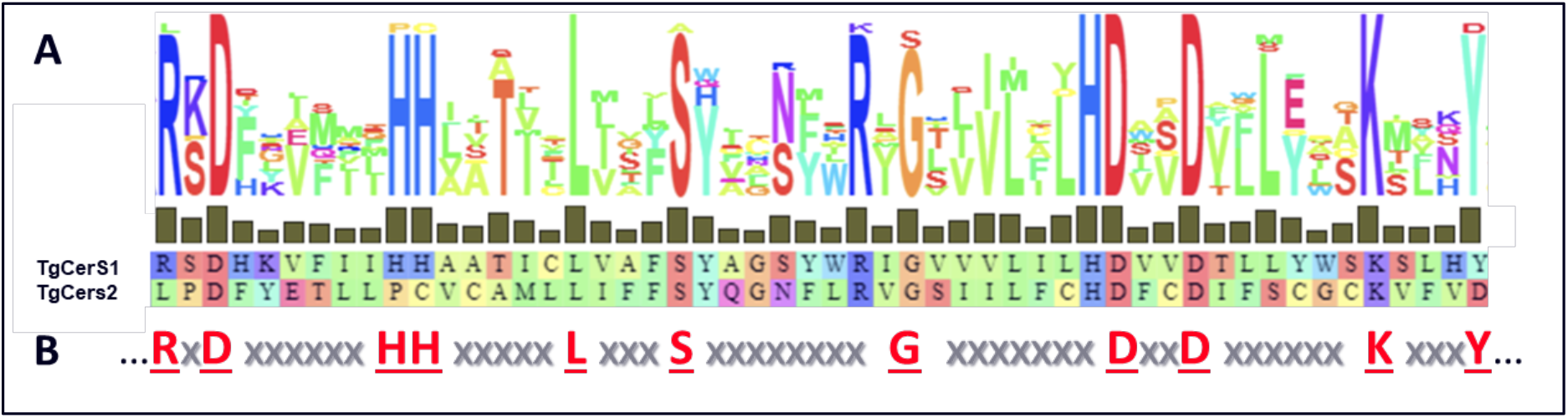
*Tg*CerS1 and *Tg*CerS2 catalytic site residues. (A) Alignment of *Tg*CerS1 (residue 208-259) and *Tg*CerS2 (residue 169-220) putative active site residues with conservation in relation to the lag1p consensus motif shown above. Hydrophobic amino acids are depicted in green; aromatic amino acids in cyan; aliphatic amino acids in red and orange; and large polar acids in purple and blue [61]. (B) Lag1p consensus motif, canonical residues depicted in red.

An initial search of the *Toxoplasma* genome database (toxodb.org/) revealed a single, putatively encoded 383 amino acid CerS orthologue, TGGT1_316450 (named *Tg*CerS1). However, re-probing the database with the identified sequence yielded an isoform, TGGT1_283710 (named *Tg*CerS2) a predicted 342 long amino acid protein which exhibited only 16.8% sequence identity and 34.5% similarity to *Tg*CerS1. *Tg*CerS2 demonstrated higher homology within the lag1p motif region of its isoform (25% identity and 42.3% similarity, CLUSTALW2 and MAFFT alignment), however it lacked the essential and canonical arginine (R208), double histidine (H217 H218) and tyrosine (Y259) residues (Figure 2A; *Tg*CerS1 numbering). All of these are subject to non-conservative changes in *Tg*CerS2, leucine (L169), proline (P178), cysteine (C179) and aspartic acid (D220) respectively (Figure 2A; *Tg*CerS2 numbering). Whilst the structure-function relationships of CerS’s are not well studied, their topology displays a conserved consensus of seven predicted transmembrane domains (TMDs) with the active site facing the lumen of the ER [44, 45] and their acyl-chain preference determined by a region of 11 residues forming an *exo*-membranous loop [46]. *Tg*CerS1 and *Tg*CerS2 maintain the consensus of seven TMDs as predicted by the computational protein structure prediction software AlphaFold2 [47] (Figure 3A and B) and, increasing the confidence in the prediction, RosettaFold [48] (Figure S1).

**Fig. 3.**
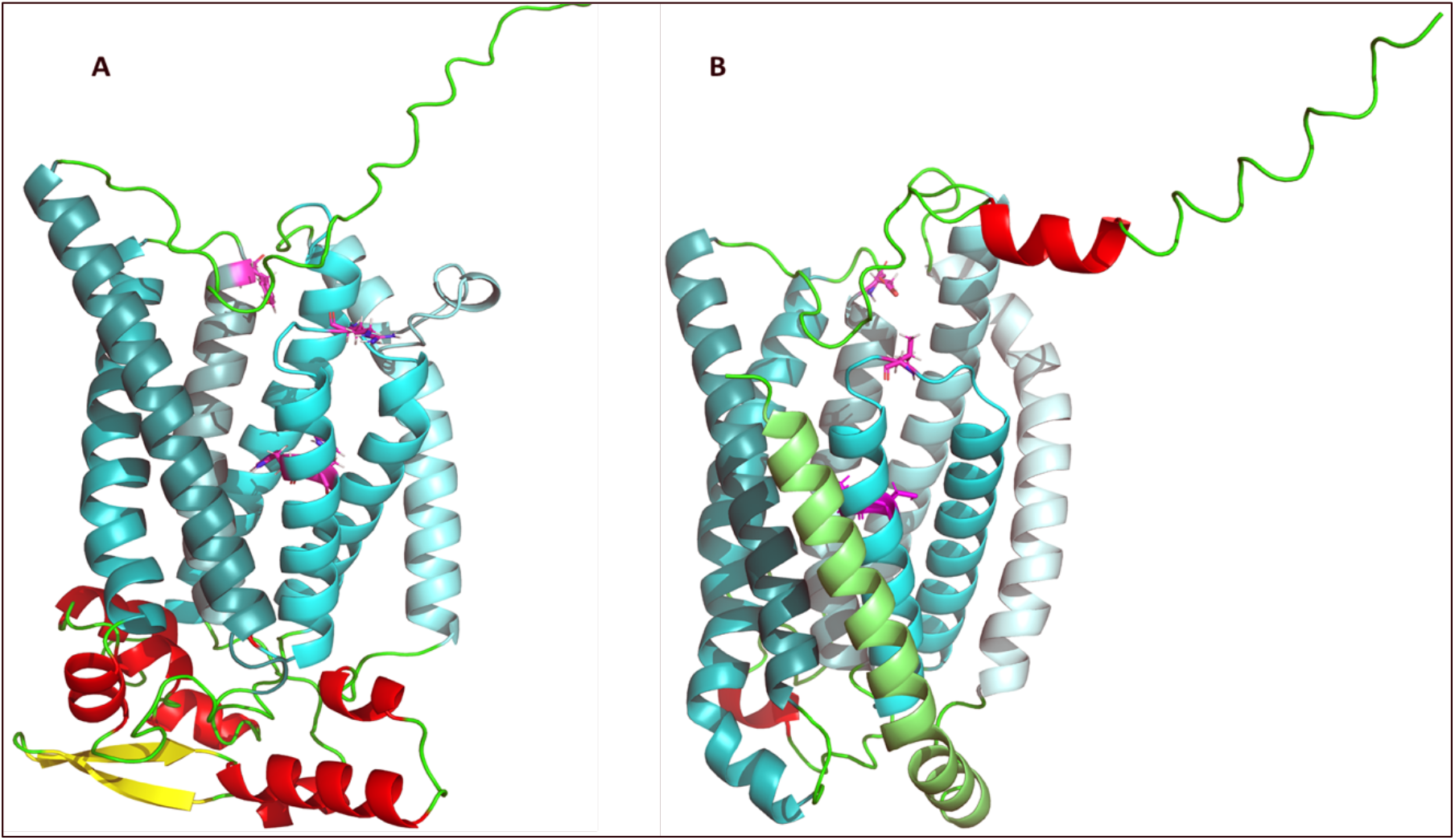
Ribbon diagram of the *ab-initio* models derived with Alphafold2 [47] of ceramide synthases (A) *Tg*CerS1 and (B) *Tg*CerS2. The transmembrane helixes are shown different shades of cyan from H1 light to H7 dark. Arginine (R208), double histidine (H217 H218) and tyrosine (Y259) residues in *Tg*CerS1, as well as leucine (L168), proline (P178), cysteine (C179) and aspartic acid (D220) residues in *Tg*CerS2, are highlighted in magenta and shown in stick formation. Putative extracellular helices shown in red, strands in yellow and loops in green. In both cases the C-termini point upwards, towards the cytosol.

The CRISPR/Cas9-based genome-wide data generated by Sidik *et al.,* 2016 [49] indicated that both orthologues have a role in parasite fitness and expression profiling demonstrated their presence throughout the lifecycle [50]. In addition, utilising a spatial proteomic approach, Hyperplexed Localisation of Organelle Proteins by Isotope Tagging (hyperLOPIT), Barylyuk *et al.*, 2020 [51] indicated that like all studied ceramide synthases, *Tg*CerS2 is located in the ER. However, the localisation of *Tg*CerS1 was not determined (Table 1).

**Table 1.** Database information regarding the identified CerS orthologues *Tg*CerS1 (TGGT1_316450) and *Tg*CerS2 (TGGT1_283710). Phenotype scores from Sidik *et al.,* 2016 [49], expression profiles from Fritz *et al.*, 2012 [50] and hyperLOPIT data from Barylyuk *et al.*, 2020 [51]. ND: Not Determined.

| Gene ID | Phenotype Score | Length (aa) | Expression | LOPIT |
| --- | --- | --- | --- | --- |
| TGGT1_316450 (*TgCerS1*) | -1.55 | 383 | Constitutive | ND |
| TGGT1_283710 (*TgCerS2*) | -1.09 | 342 | Constitutive | ER |

### Activity and localisation of *Tg*CerS1 and *Tg*CerS2

The *in vitro* study of integral membrane proteins is generally challenging. By utilising a wheat germ-based cell free membrane protein expression system, both *Tg*CerS1 and *Tg*CerS2 were expressed as FLAG-tagged proteins in *in vitro* proteoliposomes as described in Materials and Methods. Western blot analyses utilising anti-FLAG rabbit monoclonal antibodies demonstrated that both proteins are expressed (Figure 4A and Figure S2A). Subsequently, biochemical analyses of the CerS activity within these proteoliposomes was undertaken utilising fluorescent NBD-sphinganine (NBD-Sph) and acyl-CoA (C16:0 or C24:0). Lipidomic analyses have indicated that the primary ceramide species in *Toxoplasma* are C-16 and C-18 [3, 9]. In line with this observation, *Tg*CerS1 demonstrated a robust CerS activity with NBD-Sph and C16:0 acyl-CoA as substrates. No activity was observed towards NBD-Sph and C24:0 acyl-CoA (Figure 4B and Figure S2B and C). In sharp contrast, *Tg*CerS2 showed no activity to NBD-Sph combined with either of the CoA-substrates (Figure 4B and Figure S2B and C), in keeping with the absence of the canonical residues in the Lag1 motif (Figure 2A and B). The human orthologues *Hs*CerS2 and *Hs*CerS5 were analysed in the same manner and used as positive controls for CerS activity with C24:0 and C16:0 acyl-CoA respectively (Figure S2).

**Fig, 4.**
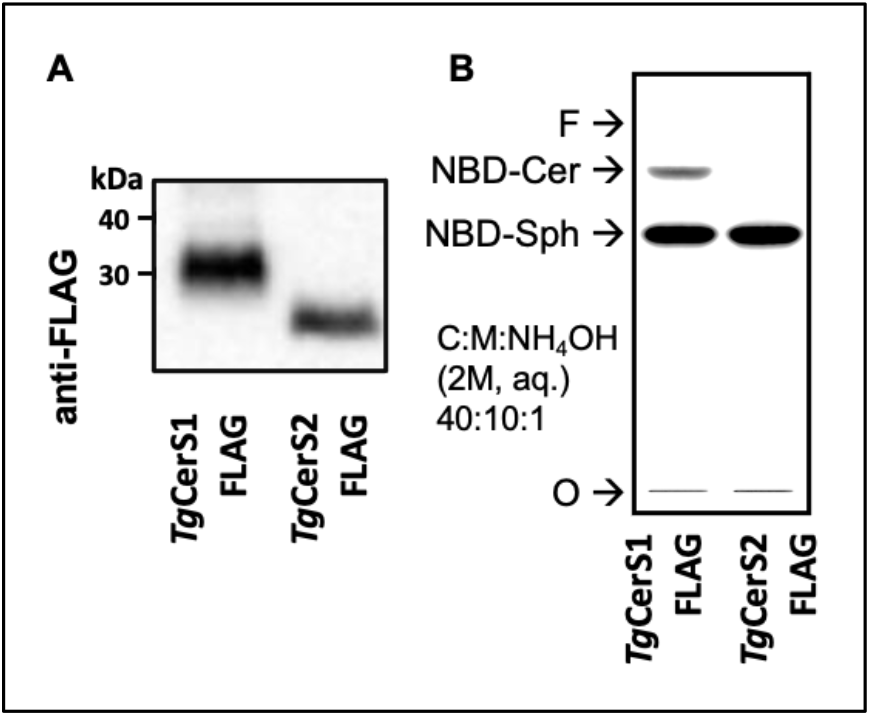
Cell-free expression and *in vitro* analyses of *Tg*CerS1 and *Tg*CerS2 (A) Western blot confirming expression of *Tg*CerS1_FLAG & *Tg*CerS2_FLAG; (B) Thin Layer Chromatography of assay products from expressed proteins using fluorescent NBD-sphinganine and C16:0 acyl-CoA as substrates; F: front, NBD-Cer: NBD-ceramide, NBD-Sph: NBD-sphinganine, O: Origin.

Like the active *Tg*CerS1, and other eukaryotic orthologues, the ‘pseudo’ ceramide synthase *Tg*CerS2 localised to ER structures in *Toxoplasma* (Figure 3A and B) when expressed from the ToxoXpress plasmid (Figure S3). As with *Tg*SPT [32], these localisations are canonical and provide further evidence that, despite the often unusual nature of the enzymes, the geometry of the sphingolipid biosynthetic pathway is conserved with respect to that seen in other eukaryotes.

### Assessment of the roles of *Tg*CerS1 and *Tg*CerS2 in parasite fitness

CRISPR/Cas9-directed knockout allowed isolation of viable, but disabled, clones lacking the *Tg*CerS1 locus. However, equivalent *Tg*CerS2 knockout clones could not be isolated. Therefore, we developed an innovative version of the conditional diCre-System [52] in which the successful excision of loxP-flanked DNA sequences is shown by a red to yellow fluorescence shift in the engineered *Toxoplasma* (Figure 6A and B). Using this system, we demonstrated that catalytically inactive *Tg*CerS2 played an even greater role in parasite proliferation and, subsequently, host cell lysis than the active orthologue *Tg*CerS1 (Figure 6C).

**Fig. 5.**
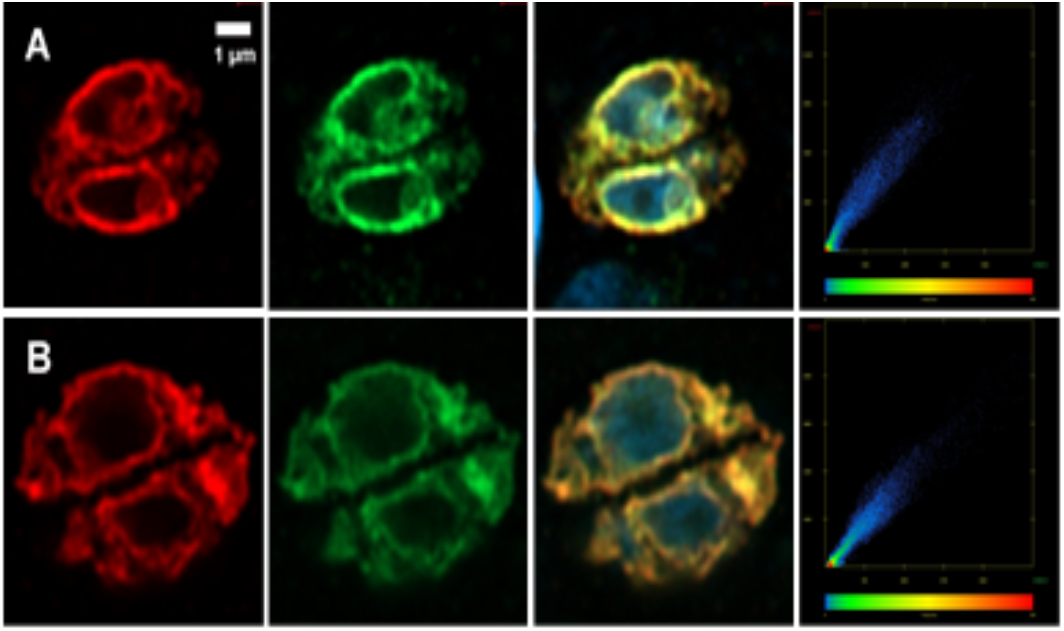
Airyscan microscopy localisation of *Tg*CerS1 and *Tg*CerS2 (A) *Tg*CerS1_FLAG and (B) *Tg*CerS2_FLAG (anti-FLAG red) relative to the ER (anti-*Tg*SPT1 green). Blue is DNA in overlay. Scatter plots (x: ER, *Tg*SPT1 green; y: (A) *Tg*CerS1 and (B) *Tg*CerS2 show overlap in signal – Pearson Correlation Coefficient: *Tg*CerS1 = 0.955 *Tg*CerS2 = 0.963 confirming that both localise to the ER.

**Fig. 6.**
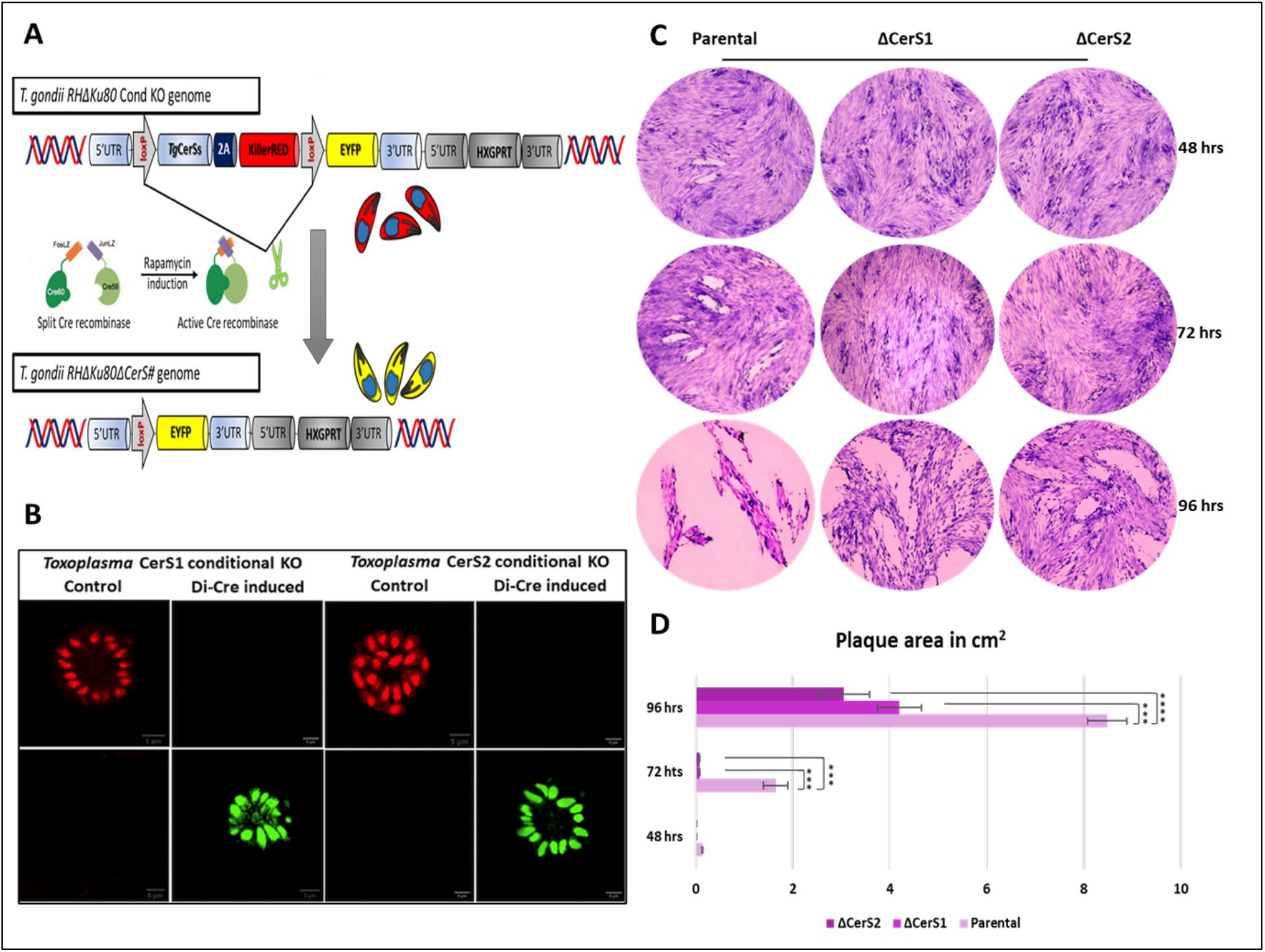
Role of *Tg*CerS1 and *Tg*CerS2 in parasite fitness (A) Schematic representation of the diCre mode of action and formation of *Tg*CerS1 and *Tg*CerS2 conditional KOs. (B) Fluorescence identification of successful *Tg*CerS1 and *Tg*CerS2 deletion after 100nM rapamycin induction. Pictures taken 72 hours post infection, scale bars 5µM. (C) Fitness phenotype of *Toxoplasma* parental and rapamycin induced *Tg*CerS1 (ΔCerS1) and *Tg*CerS2 (ΔCerS2) knockouts at 48, 72 and 96 hours post infection. Loss of either open reading frame led to significant growth impairment. (D) Numeric depiction of *Toxoplasma* plaque sizes in cm^2^, P value significance thresholds were set at: *** p<0.001, **** p<0.0001.

Specifically, plaques covered 8.48 ± 0.41 cm^2^ of the total 9.6 cm^2^ 96 hours post infection in the case of the parental *Toxoplasma* strain, compared to 4.1 ± 0.45 cm^2^ and 3.05 ± 0.52 for rapamycin induced *Tg*CerS1 and *Tg*CerS2 knockouts (Figure 6D), signifying an approximate 50% and 65% decrease in parasite mediated lysis respectively. As anticipated, uninduced *Tg*diCre:CerS1 and *Tg*diCre:CerS2 mutant lines showed the same fitness phenotype as parental (Figure S4). Notably, the non-conditional Δ*Tg*CerS1 clonal line demonstrated the same loss in fitness as its inducible equivalent under the same conditions, 4.2 ± 0.41 cm^2^ of plaques, an approximate 50% decrease compared to parental (Figure S5).

### Phylogenetic analyses of the apicomplexan ceramide synthases

Further analyses revealed that *Tg*CerS1 and *Tg*CerS2 have orthologues in the Apicomplexa. Two in *Cryptosporidium muris*, *Cystoisospora suis, Cyclospora cayetanensis, Hammodia hammondi, Plasmodium falciparum* and *Sarcocystis neurona,* but only one copy in *Cytauxzoon felis, Eimeria tenella, Neospora canium* and *Theileria annulata*. CerS amino acid sequences, including apicomplexan, human and *Arabidopsis thaliana*, were analysed using phylogenetic analyses, including Neighbour Joining (Figure 7A), Maximum Parsimony and Minimum Evolution (Figure S6). In all cases, the sequences separated into two distinct and distant clades. The first is characterised by the conserved double histidine residues (H217 H218 in *Tg*CerS1) and represents the majority of the ceramide synthases, including the subgroups of *Hs*CerS1-6, *At*CerS1-3 and the apicomplexan orthologues of the functional ceramide synthase, *Tg*CerS1. The second clade contains the ‘pseudo’ ceramide synthase, *Tg*CerS2, and its apicomplexan orthologues. All of these possess only one (Q209 H210, e.g. *Plasmodium* spp.) or no histidines (P178 C179, e.g. *Tg*CerS2) in place of the pair found in the other clade (Figure 7B). Furthermore, the R to L and Y to D changes found in the lag1 domain of *Tg*CerS2 (Figure 2; at positions 169 and 220 respectively) are conserved in all members of this clade. Coupled with the evidence presented above showing that *Tg*CerS2 is not catalytical active *in vitro*, although still ER localised, this strongly suggested that the genes encoding this apicomplexan unique ‘pseudo’ ceramide synthase diverged early in evolution probably as a result of gene duplication, where one copy of the gene maintained the original function and the other diverged to a new role. The majority of species that possess a single CerS maintain an orthologue of the functional ceramide synthase *Tg*CerS1. Assuming an early gene duplication event led to the CerS2 orthologues, this indicated that the second copy was lost during evolution. However, surprisingly *E. tenella,* a coccidian apicomplexan like *Toxoplasma*, only maintains an orthologue of the ‘pseudo’ ceramide synthase, *Tg*CerS2, indicating that this parasite is unable to synthesize ceramide *de novo*.

**Fig. 7.**
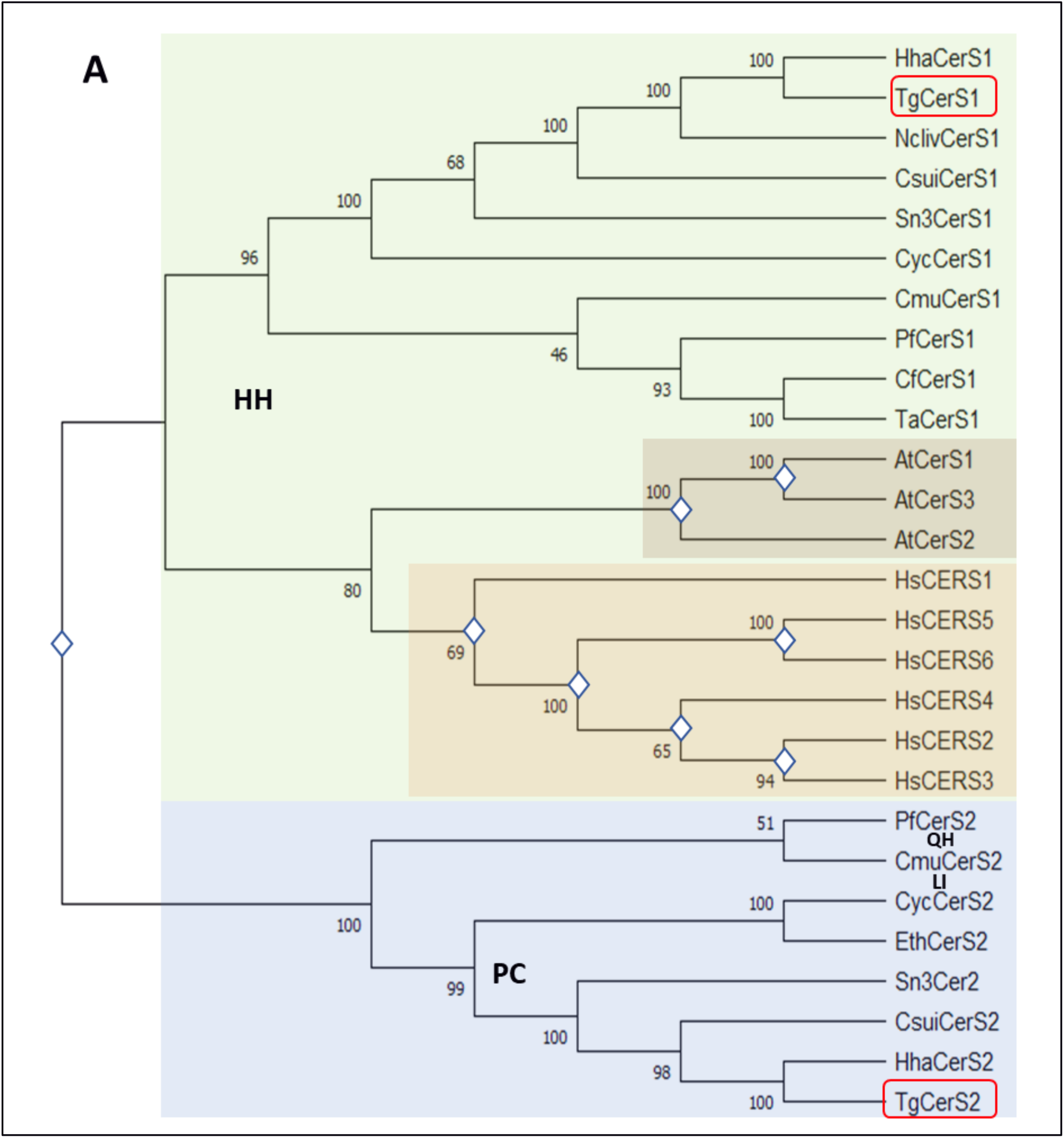
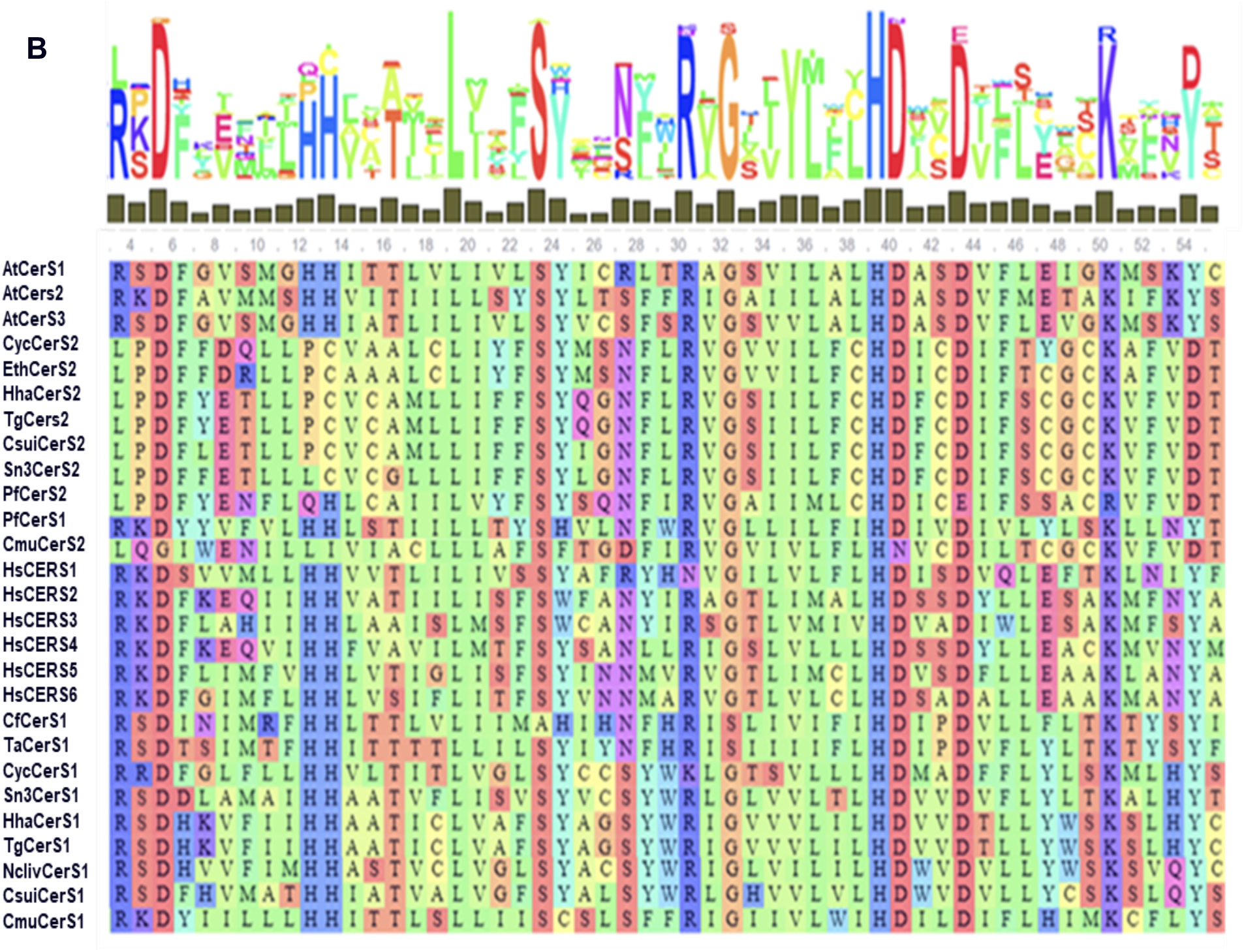
Phylogentic analyses of the apicomplexan ceramide synthases. (A) Neighbour Joining (N-J) tree of apicomplexan, human and *A. thaliana* ceramide synthases analogues. Bootstrap values are based on 1000 replicates. *Toxoplasma Tg*CerS1 and *Tg*CerS2 are highlighted in red boxes; Gene duplication assessment based on N-J tree analysis and represented by rhombuses. The distinct clades containing *Tg*CerS1 and *Tg*CerS2 are highlighted with green and purple respectively. (B) MAFFT lag1p motif alignment of *Tg*CerS1, *Tg*CerS2 and orthologues from the Apicomplexa, humans and plants. Consensus highlighted above. Hydrophobic amino acids are depicted in green, aromatic amino acids in cyan, aliphatic amino acids in red and orange, and large polar acids in purple and blue [61].

## Discussion

Due to the essentiality of sphingolipids [34], ceramide synthases (CerS) have been investigated as potential targets for therapeutic intervention in human disorders [53, 54] and microbial infections [55]. However, functional analyses have been lacking in the Apicomplexa. Therefore, in this study we sought to isolate, analyse and functionally define CerS in the model apicomplexan *Toxoplasma gondii*.

Two CerS orthologues, which we named *Tg*CerS1 and *Tg*CerS2, were identified in the *Toxoplasma* genome using bioinformatic approaches. Previously, genome-wide, analyses had identified both of these as constitutively expressed throughout the lifecycle [50] and important for parasite fitness. Here, we define the function, localisation and evolutionary origin of these unusual enzymes in the parasite. *Tg*CerS1 found to be a fully functional ceramide synthase, with preferential use of C16:0 acyl-CoA consistent with previous lipidomic analyses [3, 9]. Surprisingly, *Tg*CerS2 was catalytically inactive *in vitro* (Figure 4) and contained non-conservative substitutions in the canonical residues of the defining Lag1p motif. Utilising the recently released AlphaFold2/RosettaFold artificial intelligence programs [47, 56] we established that the structure of both orthologues was canonical, seven transmembrane domains with the substituted residues found in same arrangement in both *Tg*CerS1 and *Tg*CerS2 (Figure 3). Furthermore, using an immunofluorescent approach both proteins were identified in the *Toxoplasma* ER, the canonical localisation of CerS (Figure 5). Together, these data indicated that *Tg*CerS2, whilst non-catalytic, had maintained structure, localisation and, presumably, functionality through evolution. Furthermore, using an inducible knockout approach, both orthologues were shown to have an important role in tachyzoite proliferation with the loss of the non-catalytically active *Tg*CerS2 demonstrating an even greater effect than ablation of *Tg*CerS1 (Figure 6).

From where did this ‘pseudo’ enzyme arise? Further bioinformatic analyses demonstrated that orthologues of *Tg*CerS1 could be identified in almost all members of the Apicomplexa, and *Tg*CerS2 orthologues were found in a large subset. Phylogenetic analyses (Figure 7A and S6) clearly demonstrated that the predicted ceramide synthases divide into two clades, one containing the *Tg*CerS1 orthologues and including the *Arabidopsis* and human enzymes, the other clustered the *Tg*CerS2 ‘pseudo’ enzyme orthologues from the Apicomplexa phylum. Gene duplication analyses (Figure 7A) indicated that this occurred early in evolution for the Apicomplexa, prior to speciation. The fact that all the *Tg*CerS2 orthologues maintain the R-L and Y-D changes (positions 169 and 220 in *Tg*CerS2) and the loss of canonical HH (positions 217 and 218 in *Tg*CerS2), to PC in all aside from *Plasmodium* spp. (QH), supported early divergence and the maintenance of key functionality. However, despite this putative selective pressure, *Cytauxzoon felis, Theileria annulata* and *Neospora canium* encode only a *Tg*CerS1 orthologue (Figure 7B). Whilst *C. felis* and *T. annulata* are of the same family, Theileriidae, *N. canium* is in the Sarocystidae. *Sarcocystis neurona, Cystoisospora suis* and *Hammodia hammondi* are also in this latter family and maintain orthologues of both *Tg*CerS1 and *Tg*CerS2 (*T. gondii* is also in the Sarocystidae). This suggested that the loss of the *Tg*CerS2 orthologue occurred later in evolution due to unknown environmental stresses and evolutionary pressures.

In contrast, and interestingly, *E. tenella* maintain a *Tg*CerS2 orthologue but appears to have lost the active ceramide synthase which indicated that this parasite lacks the ability to synthesize ceramide *de novo* (Figure 7). This further supported the importance of the non-catalytic ceramide synthase, a hypothesis reinforced by the same gene arrangement being found other *Eimeria* species (see Materials and Methods). However, uniquely in the *Eimeria* species, *E. falciformisi* maintains the functional *Tg*CerS1 and ‘pseudo’ *Tg*CerS2 orthologue arrangement (EfaB_MINUS_15758.g1433 and EfaB_PLUS_2387.g283 respectively). Interestingly, *E. falciformisi* is the only *Eimeria* species in which the sphingolipid content has been extensively analysed, with a lipidomic approach indicating that *de novo* ceramide biosynthesis occurs [10]. Like *E. falciformisi,* another member of the family Eimeridae, *Cyclospora cayetanensis,* also encoded ortholgues for both proteins. Together, these data indicated that loss of orthologues of either protein (*Tg*CerS1 or *Tg*CerS2) is a relatively recent event in evolution and that at least one must be maintained. Apart from in most *Eimeria* species, the maintained orthologue was a catalytical active ceramide synthase.

However, the maintenance of apicomplexan *Tg*CerS2 orthologues, either alongside a functional enzyme or, in the *Eimeria* species, alone is mysterious given its lack of catalytic activity. Following gene duplication, one copy could become non-functional due to degenerative mutations. However, alongside the situation in *Eimeria* species, the conservation of amino acids in the Lag1 motif and of the tertiary structure (Figures 2 and 3) in these orthologues make this hypothesis unlikely. Alternatively, one of the duplicated copies could acquire a new function that is favourable for the organism. The maintenance of *Tg*CerS2 orthologues, and its non-catalytic nature, across the Apicomplexa support this hypothesis. However, the true function of the *Tg*CerS2 orthologues remains unknown. Given the proposed dimerization of human CerS2 and CerS5 to modulate activity [57] we investigated the possibility of *Tg*CerS1 and *Tg*CerS2 dimerization using the predicted structures (Figure 3) and those similarly generated for human CerS2 and CerS5. However, no evidence for the formation of dimers was provided for either pair (data not shown), although apparent *Tg*CerS1 and *Tg*CerS2 homodimers were seen by Western blotting (Figure S2A). In addition, the conservation of *Tg*CerS2 alone in the *Eimeria* spp. renders this explanation unlikely. An alternative hypothesis would be that the non-catalytic isoforms act as ceramide binders and regulate the availability of this highly bioactive lipid. This is a particularly attractive explanation for the *Eimeria* species which presumably would have to scavenge and regulate host ceramide.

Whilst functional analyses of these essential proteins in an obligate intracellular pathogen of a host encoding the same activity is highly challenging, further detailed analyses are required to fully understand the roles of the apicomplexan ‘pseudo’ enzyme. However, the identification and analyses described here detail a system unique to the Apicomplexa, a phylum of parasitic protozoa. Given the importance to parasite proliferation, coupled with the non-mammalian nature of ceramide synthesis and regulation, this could represent a potential point for therapeutic intervention for a range of diseases from malaria to toxoplasmosis.

## Materials and Methods

### Culturing of *Toxoplasma gondii* and host cells

Human foreskin fibroblasts (HFFs) (SRC-1041, ATCC®) were cultivated in culture treated plastics (T-75s; T-25s and 6 well plates) presence of Dulbecco’s modified Eagle’s medium (DMEM-Gibco) supplemented with 10% foetal bovine serum (SIGMA), 2mM L-glutamine and 1% penicillin-streptomycin solution. HFF cells were not used beyond passage 20. All strains of *Toxoplasma gondii* including RH.diCre.Δku80, RH.ΔKu80.ΔCerS1, RH.diCre.Δku80:CerS1 and RH.diCre.Δku80:CerS2 were maintained *in vitro* by serial passages on monolayers of HFFs maintained at 37°C, 5% CO_2_ in a humidified incubator. Freshly egressed parasites were assessed for viability by Trypan blue (0.4% w/v) to ensure high viability before downstream experiments.

### Isolation of *Toxoplasma* genomic DNA and RNA

Genomic DNA was extracted from *T. gondii* tachyzoites to use as a PCR template by pelleting parasites and resuspending in PBS. DNA extraction was then performed using the QIAamp DNA blood mini kit (Qiagen) as per the manufacturer’s protocol. RNA was isolated from freshly purified tachyzoites of *Toxoplasma gondii* using RNeasy Micro Kit (Qiagen) subsequently reverse-transcribed into first-strand cDNA using Qiagen one step RT-PCR kit System.

### Cell-free Expression *Tg*CerS1 and *Tg*CerS2

*Tg*CerS1 and *Tg*CerS2 open reading frames were synthesized (GenScript) following PCR-amplification using primer pairs *Tg*CERS1.CFE.P1 / *Tg*CERS1.CFE.P2_FLAG and *Tg*CERS2.CFE.P1 / *Tg*CERS2.CFE.P2_FLAG respectively (Table S3) and InFusion (TaKaRa) cloned into pEU-E01-MCS. The mammalian pCMV-Tag2B-CerS2 / CerS5 expression vectors were a kind gift from Tony Futerman, Weizmann Institute, Israel. HsCerS2-V5-His and FLAG-HsCerS5-V5-His were PCR amplified and cloned into the pEU Flexi vector (a kind gift from James Bangs, University of Wisconsin, Madison). To this end, first a KpnI restriction site was introduced directly between the hSMS1 ORF and V5 tag in the pEU-Flexi vector using site-directed mutagenesis according to the Stratagene Quick Change protocol with primers pEU.Flexi.P1 and pEU.Flexi.P2 (Table S3). Then HsCerS2-V5-His and FLAG-HsCerS5-V5-His were PCR-amplified using primer pairs *Hs*CERS2.CFE.P1 / *Hs*CERS2/5.CFE.P2 and *Hs*CERS5.CFE.P1 / *Hs*CERS2/5.CFE.P2 respectively (Table S3). The PCR products were ligated into the pJET1.2 blunt vector (Thermo Scientific), digested with XhoI and KpnI alongside with the pEU-KpnI Flexi acceptor vector, and the final constructs were obtained by T4 ligation (Invitrogen). Using the Protein Research Kit S16 (CellFree Sciences) cell-free expression of *Tg*CerS1, *Tg*CerS2, *Hs*CerS2-V5-His and FLAG-*Hs*CerS5-V5-His was performed as described for sphingomyelin synthases [58] but without the application of a dialysis reservoir and in the presence of 100 nm liposomes consisting of PC:PE:PI 2:2:1 (mol:mol:mol). Protein expression was confirmed by SDS-PAGE and Western blotting using ready-made gels (Invitrogen; 100V, 60 min) before transferring to a nitrocellulose membrane (Life Technologies; BioRad; 100V for 60 min). Following blocking (2 mM Tris-HCl, 50 mM NaCl, pH 7.5, 0.5ml Tween 20 and 5% w/v fatty-acid-free BSA 60 mins at room temperature), protein expression was determined by Western blotting using as primary antibodies anti-FLAG or anti-V5 tag antibody (AbCam; 1:500, overnight at 4°C. Rabbit anti-Mouse IgG (H+L) Secondary Antibody, HRP (ThermoFisher; 1:5000, 60 min at 4°C). Detection was achieved with Pierce™ ECL Western Blotting Substrate using a ChemDoc XR+ (BioRad).

### Ceramide Synthase Assay

Following determination of expression, equivalent concentrations of *Tg*CerS1 and *Tg*CerS2, plus controls, were assayed for ceramide synthase activity in LoBind tubes (Eppendorf) as follows. A 100 µl reaction volume with 80µl of proteoliposome mixture, 15µl cell free expression translation buffer (15 mM HEPES-KOH (pH 7.8), 50 mM potassium acetate, 1.25 mM magnesium acetate, 0.2 mM spermidine hydrochloride, 0.6 mM ATP, 0.125 mM GTP, 8 mM creatine phosphate), 1µl NBD-Sph (5mM stock; Avanti Polar Lipids Inc.), 1µl Acyl-CoA C16:0 / C24:0 (5mM stocks; Avanti Polar Lipids Inc.), 4µl delipidated BSA (5µM stock; Sigma Aldrich). Following incubation (60 min at 37°C) the reaction was halted by addition of 375µl chloroform:methanol (1:2) and lipid extracted according to Bligh and Dyer. Following drying and re-suspension in 10:10:3 chloroform:methanol:H_2_O lipid extracts were transferred to a NANO-ADAMANT HP-TLC plate (Macherey & Nagel) using an ATS5 TLC sampler (CAMAG, Berlin, Germany). The TLC was developed in CHCl_3_:MeOH:2M_aq_NH_4_OH (40:10:1 v:v:v) or CHCl_3_:MeOH:H_2_O (80:12:1 v:v:v) in a CAMAG ADC2 automatic TLC developer [59]. The NBD-lipids were detected using a Typhoon FLA 9500 biomolecular imager (GE Healthcare Life Sciences) operated in Cy2 fluorescence mode with 473 nm excitation laser, BPB1 filter, 50 μm pixel size, and PMT voltage setting of 290 V. Data were later processed using ImageLab (Bio-Rad Laboratories) to adjust intensity.

### pTXP-*Tg*CerS1-FLAG, pTXP-*Tg*CerS2-FLAG cloning and subcellular localization of *Tg*CerS1 and *Tg*CerS2

Primers were designed to amplify and FLAG tag the *Tg*CerS1 and *Tg*CerS2 coding sequences: *Tg*CerS1Xpress / *Tg*CerS1FLAGXpress_R and *Tg*CerS2Xpress_F / *Tg*CerS2FLAGXpress_R from the synthetic genes (Table S4). The resultant PCR products were introduced into NheI (NEB) linearized ToxoXpress vector (pTXP) using In-Fusion HD cloning kit (TaKaRa) as per manufacturer’s protocol (Figure S3) to create pTXP_*Tg*CerS1-FLAG and pTXP_*Tg*CerS2-FLAG. All constructs were verified by sequencing (MWG) prior to further work. Transfections were carried out using a 4D Nucleofector (Lonza), protocol FI 158. Briefly, parasites freshly lysed from HFFs monolayer were homogenized by passage through a 25-gauge needle and isolated by centrifugation at 1500 × *g* for 10 min at 4 °C. The pellet was resuspended in the P3 buffer with added supplementary buffer P1 (Lonza). 20 μl of the parasite suspension (10^7^ ml^−1^) were added to a dried pellet of ethanol-precipitated pTXP_*Tg*CerS1-FLAG or pTXP_*Tg*CerS2-FLAG plasmid and electroporated. Subsequently, 100 μl of medium was added, and 10 μl or 20 μl were added to 24-well plates containing confluent HFF cells grown on glass coverslips and the plates incubated at 37 °C, 5.0% CO_2_. Cells were fixed in 4% paraformaldehyde in PBS (pH 7.4) for 15 minutes and then permeabilized with 0.4% (v/v) Triton X-100 in PBS for 10 min, before incubation in blocking buffer (PBS supplemented with 1% (w/v) BSA; Sigma-Aldrich), 0.1% fish skin gelatin (Sigma-Aldrich), and 0.1% (v/v) Triton X-100) for 15 minutes at room temperature. Samples were incubated with a rabbit monoclonal anti-FLAG antibody (1:200; AbCam) or the primary anti-*Tg*SPT1 Δ158 rat polyclonal [32] (1:200) in blocking buffer overnight at 4 °C. After PBS washing, samples were incubated with Alexa Fluor® 594 anti-rabbit and Alexa Fluor® 488 anti-rat secondary antibodies (ThermoFisher) at 1:500 in blocking buffer for 1 hour at room temperature. The samples were incubated with DAPI (Sigma-Aldrich) in PBS for 10 min, mounted using Vectashield H-1000 (Vector labs), and sealed before imaging. Co-localization was assessed using the ScatterJn plugin and scatter plots [60] as previously [32]. A linear correlation demonstrates a strong spatial correlation between the channels, and the slope indicated the relative intensities.

### Construction of *Tg*CerS1 and *Tg*CerS2 plasmids for CRISPR/Cas-9-based manipulation

CRISPR/Cas-9 was utilised to accelerate the generation of the conditional Δ*Tg*CerS1 and Δ*Tg*CerS2 lines and also to create CRISPR/Cas-9 KOs. All CRISPR/Cas9 plasmids designed and used in this study were created by using Q5 Site-Directed Mutagenesis Kit (NEB-E0554S) based on the previously described CRISPR/Cas9 plasmid [58] as primary scaffold, as per the manufacturer’s protocol. All constructs were verified by sequencing (MWG) prior to further work. Primers for *Tg*CerS1 and *Tg*CerS2 sgRNAs were predicted and designed using online tools including <u>EuPaGDT</u>, <u>E-CRISP</u> and <u>CHOPCHOP</u> (Table S5), targeting the 5’ and 3’ of the respective coding regions to ensure efficient gene deletion and subsequent DNA insertion. CRISPR/Cas-9 plasmids were transfected along with supplementary DNA cassettes to allow mutant verification and positive selection. In the case of non-conditional knockouts, a GFP reporter gene was introduced to the corresponding genes *Tg*CerS1 and *Tg*CerS2 respectively, after DNA amplification from a previously described vector [59] using set of primers (Table S6).

### Construction of *Tg*CerS1 and *Tg*CerS2 cassettes for inducible recombinase-based knockout

A pUC19 (NEB) plasmid was used as a backbone for the constructs after it was linearized with FD PvuII to customise the subsequent assembly of the appropriate fragments. All DNA amplicons for plasmids constructs were amplified with high fidelity Phusion Polymerase (NEB) to minimize replication errors. pUC19-5’UTR-loxP-*Tg*Cers1-2A-KillerRed-loxP-EYFP-3’UTR-5’UTR-HXGPRT-3’UTR and pUC19-5’UTR-loxP-*Tg*CerS2-2A-KillerRed-loxP-EYFP-3’UTR-5’UTR-HXGPRT-3’UTR was designed to be knocked-in into the *Tg*CerS1 or *Tg*CerS2 loci respectively. Briefly, 4 different DNA fragments were amplified using primers sets F01, F02, F03 and F04 (Tables S6 and S7; Integrated DNA Technologies [IDT]). Fragments were cloned into a PvuII linearized pUC19 using In-Fusion HD cloning kit (TaKaRa) as per manufacturer’s instructions. All constructs were verified by sequencing (MWG) prior to further work. FastDigest (FD) LguI and a combination of FD SacI / FD Ajul (for *Tg*Cers1) and FD XbaI and combination of FD ScaI and FDHind III (for *Tg*Cers2) were used to digest plasmids at 37°C for 60 minutes and electrophoresed on a 0.8% agarose gel to verify successful assembly. Finally, FD SphI and FD CpoI were used to linearize CerS1 DNA cassette and FD HindIII was used to linearize CerS2 DNA cassette before electroporated into freshly egressed RH.diCre.Δku80 (parental) parasites as described above.

### *Toxoplasma* mutant strains selection and verification

All transfections were performed by a 4D Nucleofector (Lonza) as described above. Transfected parasites were incubated overnight at 37 °C and 5% CO_2_ with media being replaced with complete DMEM containing 25μg/mL MPA (Sigma-Aldrich) and 50μg/mL xanthine (Sigma-Aldrich) the following day and left to grow for at least 8 days with the selection media, refreshed every 48 hours. Clonal line selection was then performed with newly acquired mutant strains before molecular verification by diagnostic PCR using GoTaq DNA polymerase (Promega) and multiple sets of primers (Tables S9 and S10). In addition, confocal fluorescence microscopy monitoring the red (585/610nm) to yellow (513/527nm) fluorescence switch was utilised as described below. CRISPR/Cas-9 non-conditional KOs were also assessed microscopically monitoring GFP signal post transfection.

### Conditional *Tg*CerS1 and *Tg*CerS2 KΟs

Δ*Tg*CerS1 and Δ*Tg*CerS2 conditional knockouts were obtained after the addition of 100nΜ rapamycin to RH.diCre:CerS1 and RH.diCre:CerS2 strains respectively. Briefly, tachyzoites were collected from HFFs by scraping cells and passing through a 25G needle by syringe lysis. Following centrifugation at 50 x g for 5 minutes to allow removal of cell debris, the supernatant was centrifuged again at 500 x g for 5 minutes. Parasites were then resuspended and incubated for 4 hours at 37 °C, 5% CO_2_ in complete DMEM with 100nM of rapamycin before washing twice with warm media to remove rapamycin excess. Following viability assessment by trypan blue (0.4% w/v), the parasites were then used to infect HHFs monolayers. *Toxoplasma* parental line was also tested under the same treatment conditions to exclude rapamycin effect in parasite fitness.

### Confocal fluorescence microscopy

*Tg*CerS1-FLAG, *Tg*CerS2-FLAG, RH.diCre.Δku80, RH.diCre:CerS1 and RH.diCre:CerS2, un-induced and rapamycin induced parasites, were allowed to invade and replicate in HHFs monolayer grown on glass coverslips as previously described. Fluorescence images were acquired using laser scanning confocal microscope Zeiss LSM 880 with AiryScan equipped with excitation laser 405, Argon 458, 488, 514, He-Ne 543, 594, and 633 and AiryScan filter set combinations BP 420–480 + BP 495–550, BP 420–480 + BP 495–620, BP 420–480 + LP 605, BP 465–505 + LP 525, BP 495–550 + LP 570, and BP 570–620 + LP 645 (Durham Centre for Bioimaging Technology). For each image, the dynamic range was checked to avoid saturation, except with the DAPI staining where host cells masked the detection of parasite nuclei at low gain/laser power values. AiryScan images were automatically processed using default values. 40X, 63X and 100X oil lenses were used to capture red and yellow/green fluorescence. CZi outputs were exported and analysed using primarily ImageJ and Imaris software.

### Phenotypic characterization – *Toxoplasma* fitness - plaque assays

HHFs cells were grown to confluence in 6 well plates as previously described. 10^5^ parasites per well, including parental and conditional knockouts, were inoculated and incubated for 48, 72 and 96 hrs, after which wells were washed twice with PBS and fixed with 4% paraformaldehyde (pH=7.4) for 20 mins at room temperature. 0.05% w/v Crystal Violet (CV) solution (Sigma-Aldrich) was later applied to the monolayer and left for 30-40 mins at room temperature. CV solution was later removed, wells were washed twice with 2-3 ml of distilled water and left approximately two hours at room temperature to dry out. Wells were then coated with 0.3% agarose and left to dry out for 30 minutes at room temperature. Plaques were photographed using an Olympus inverted microscope CKX53 and a 5 MP XCAM camera 1080p, under 4X magnification. ImageJ was then used to determine plaques sizes utilizing the plugins Level Sets and Find Edges. Values are expressed as mean values of three independent experiments ± SD in 48, 72 and 96 hours post infection. P-values were calculated using unpaired two-tailed Student’s t-test unless stated otherwise. P value significance thresholds were set at: * p<0.05, **p<0.01, *** p<0.001, **** p<0.0001. All significant results are mentioned with asterisks in the graphs.

### Phylogenetic analysis

<u>ToxoDB</u>, <u>VEuPathBD</u>, <u>TriTrypBD</u>, <u>PlasmoDB</u>, <u>Uniprot</u>, were used in order to identify CerS analogues in various taxa and species. All accession numbers listed in the text aside from: *E. necatrix* ENH_00074610.1-p1, *E. maxima* EMWEY_00008780-t26_1-p1, *E. mitis* EMH_0037690-t26_1-p1, *E. acervuline* EAH_00007580-t26_1-p1 and *E. brunetti* EBH_0045800-t26_1-p1). Multiple amino-acid alignment was performed by using MAFFT and MUSCLE alignment algorithms. Molecular Evolutionary Genetics Analysis (MEGA-X) was used to reconstruct phylogenetic trees and perform gene duplication analysis. Neighbour Joining, UPGMA Tree and Maximum Likelihood methods were used to allow for equal and unequal rates of evolution between the CerS’s of different species and taxa. NGphylogeny.fr was also used to visualise MAFFT alignments and identify CerS consensus regions. Sequence accession numbers with respective organism names are described in Table S2.

## Acknowledgments

We thank Professor Ariel Silber (University of Sao Paulo) for helpful discussions regarding metabolism and enzyme analyses. We also thank Dr Tim Hawkins (Durham University) for support with the microscopy.

## Supporting Information

**Fig. S1.**
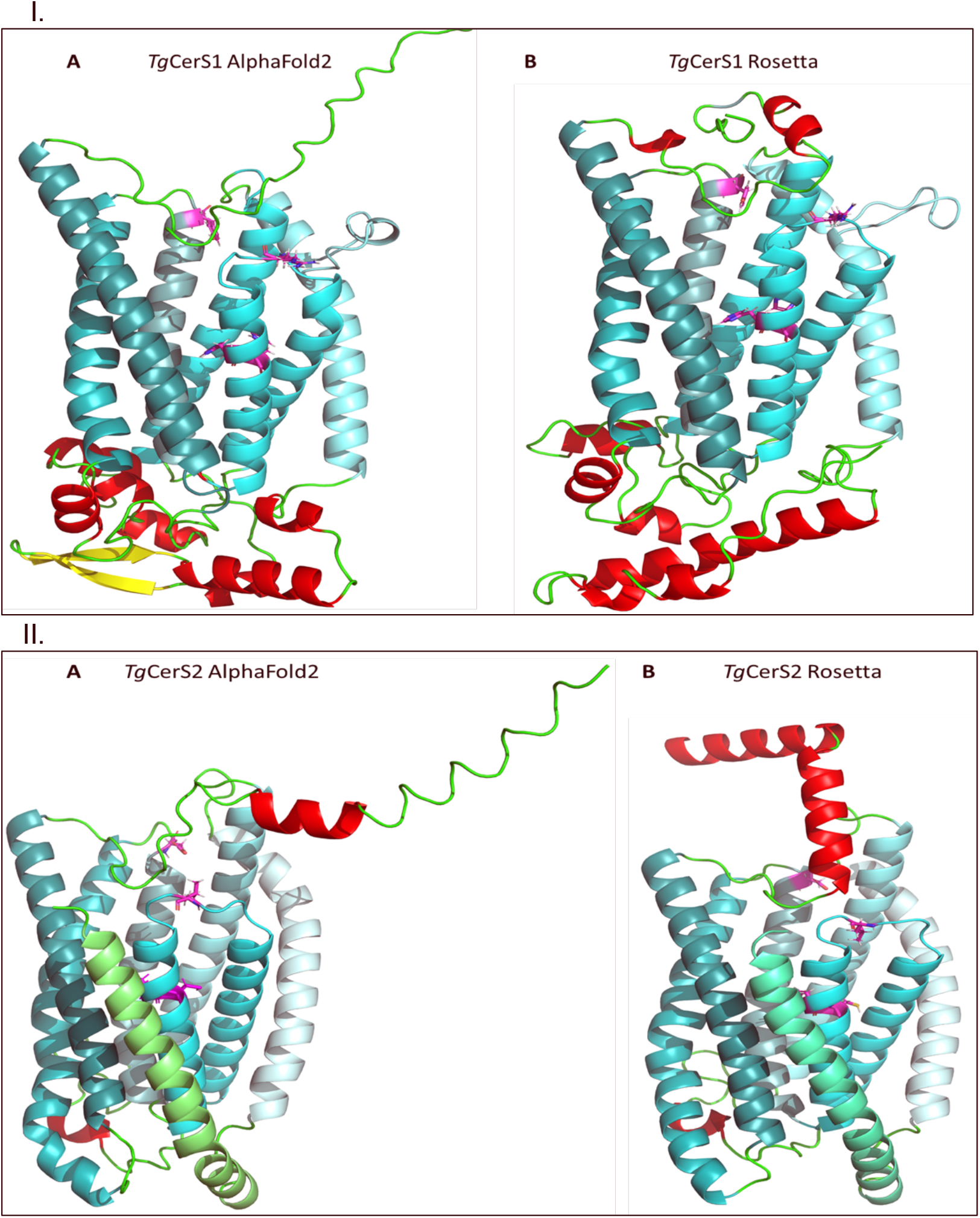
Ribbon diagram of I. *Tg*CerS1 and II. *Tg*CerS2 as predicted by (A) AlphaFold2 [47] and (B) Rosetta fold [48]. Transmembrane domains are shown in different shades of cyan from H1 - light to H7 dark. Arginine (R208), double histidine (H217 H218 and tyrosine (Y259) residues in *Tg*CerS1, as well as leucine (L168), proline (P178), cysteine (C179) and aspartic acid (D220) residues in *Tg*CerS2, are highlighted in magenta and shown in stick formation. Putative extracellular helices shown in red, strands in yellow and loops in green. C-termini point up towards cytosol.

**Fig. S2.**
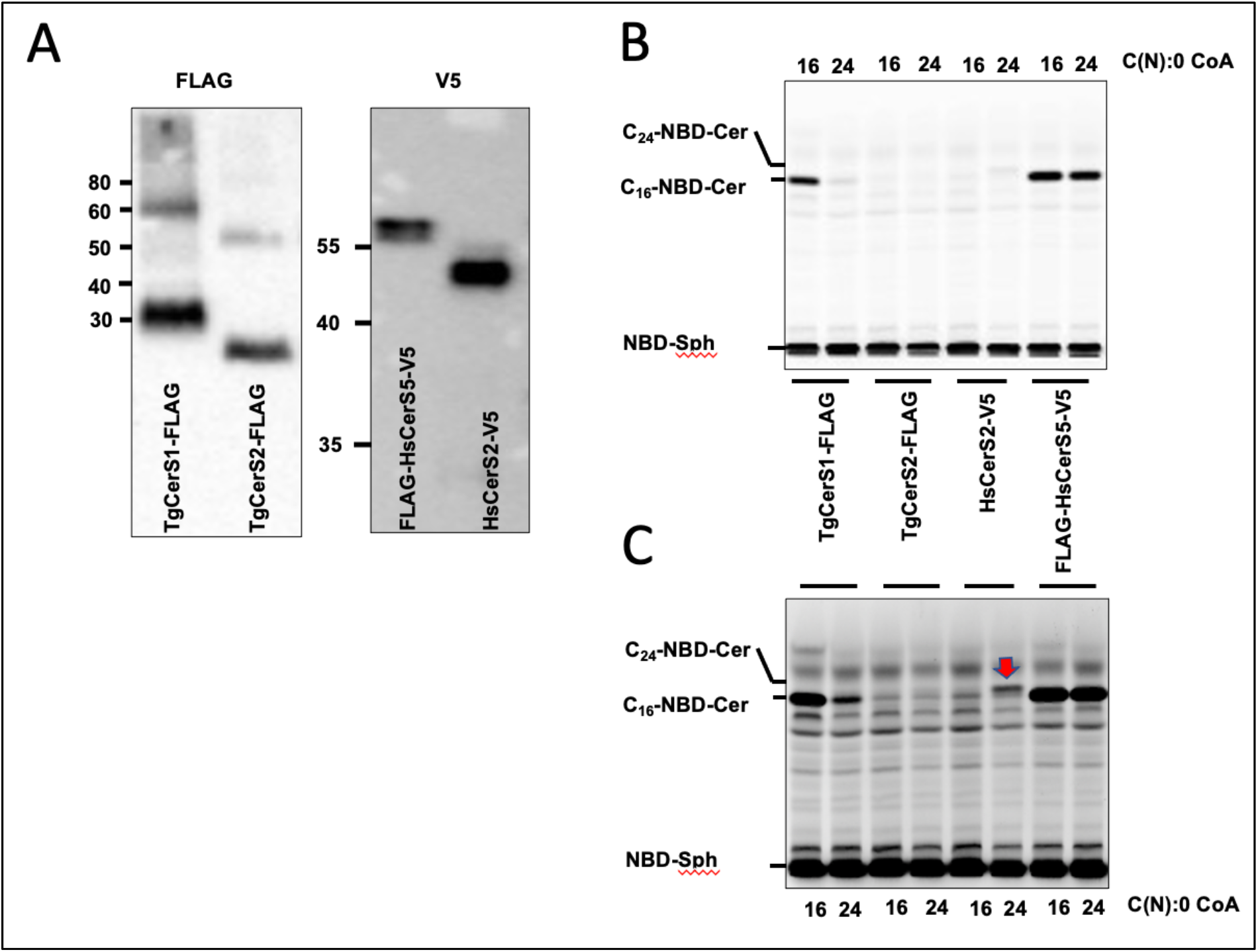
Cell-free expression and biochemical analyses of *Tg*Cer1 and *Tg*Cer2. **(A)** Western blot of expressed FLAG-tagged *Tg*Cer1 and *Tg*Cer2 (expanded from Figure 4) and V5-tagged *Hs*CerS5 and *Hs*CerS2 probed with the primary antibodies indicated. **(B)** TLC separation of ceramide synthase reaction products in solvent phase CHCl3:MeOH:H2O (80:12:1 v:v:v), *Hs*CerS5 and *Hs*CerS2 are controls and provide markers for sphinganine (NBD-Sph) and ceramide (C16-NBD-Cer and C24-NBD-Cer **respectively**). Acyl-CoA substrate for each reaction shown along top of the plate. The formation of C16-NBD-Cer by CerS5 and *Tg*CerS1 is only partially dependent on 16:0 CoA addition, presumably because the reaction mixture contains some wheat-germ derived CoA. **(C)** Longer exposure of B, red arrow indicates C24-NBD-Cer produced through action of *Hs*CerS2. The relative low activity of *Hs*CerS2 is consistent with our laboratory experience and maybe due to a lack of phosphorylation [60] or low *in vitro* bio-availability of the long chain CoA. The acyl-CoA substrate for each reaction shown along bottom of the plate.

**Fig. S3.**
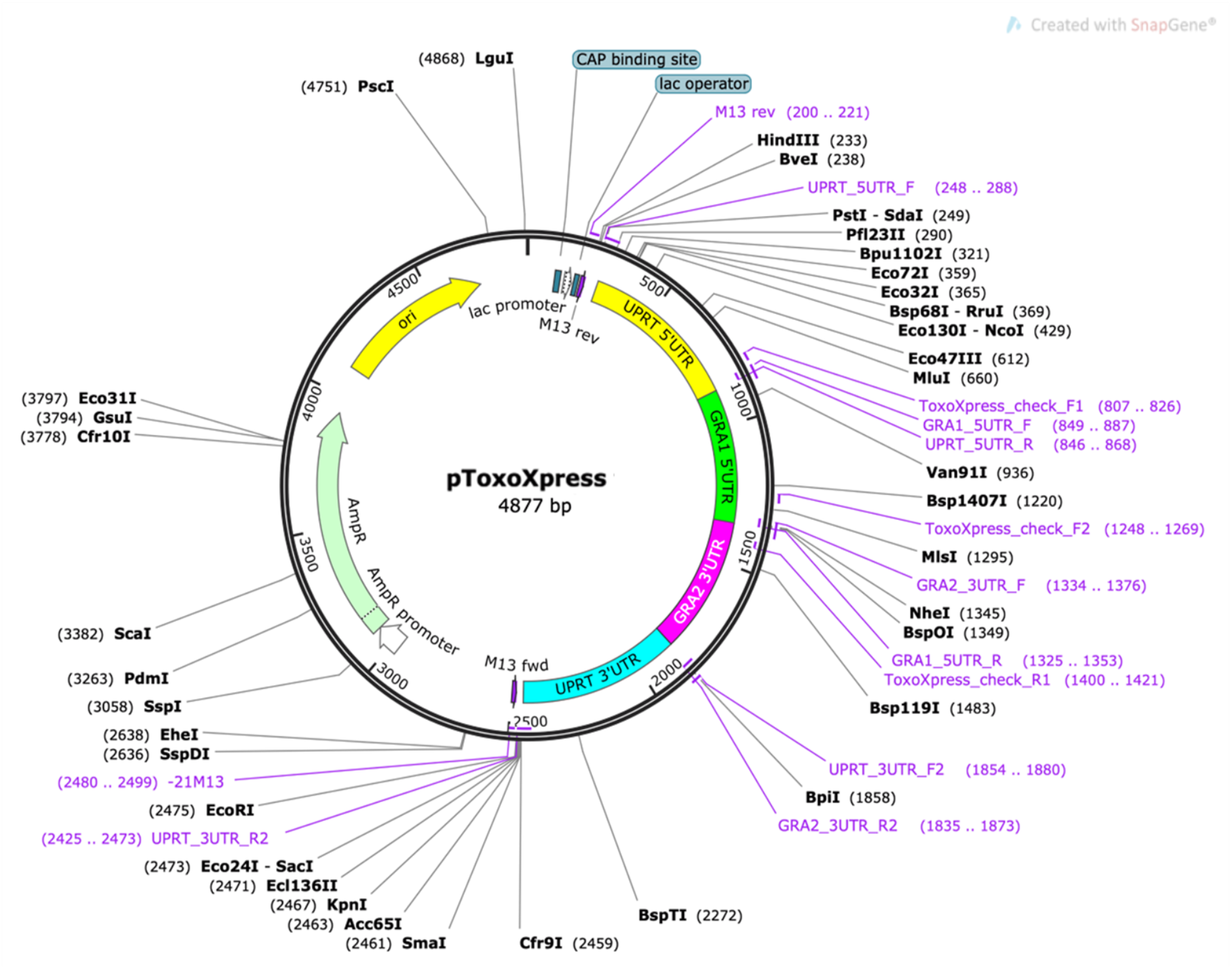
ToxoXpress was designed, created and validated in-house for either transient expression or integration into the UPRT locus and selection using 5-fluoro-2’deoxyuridine. In both cases expression is driven by the GRA1 promotor.

**Fig. S4.**
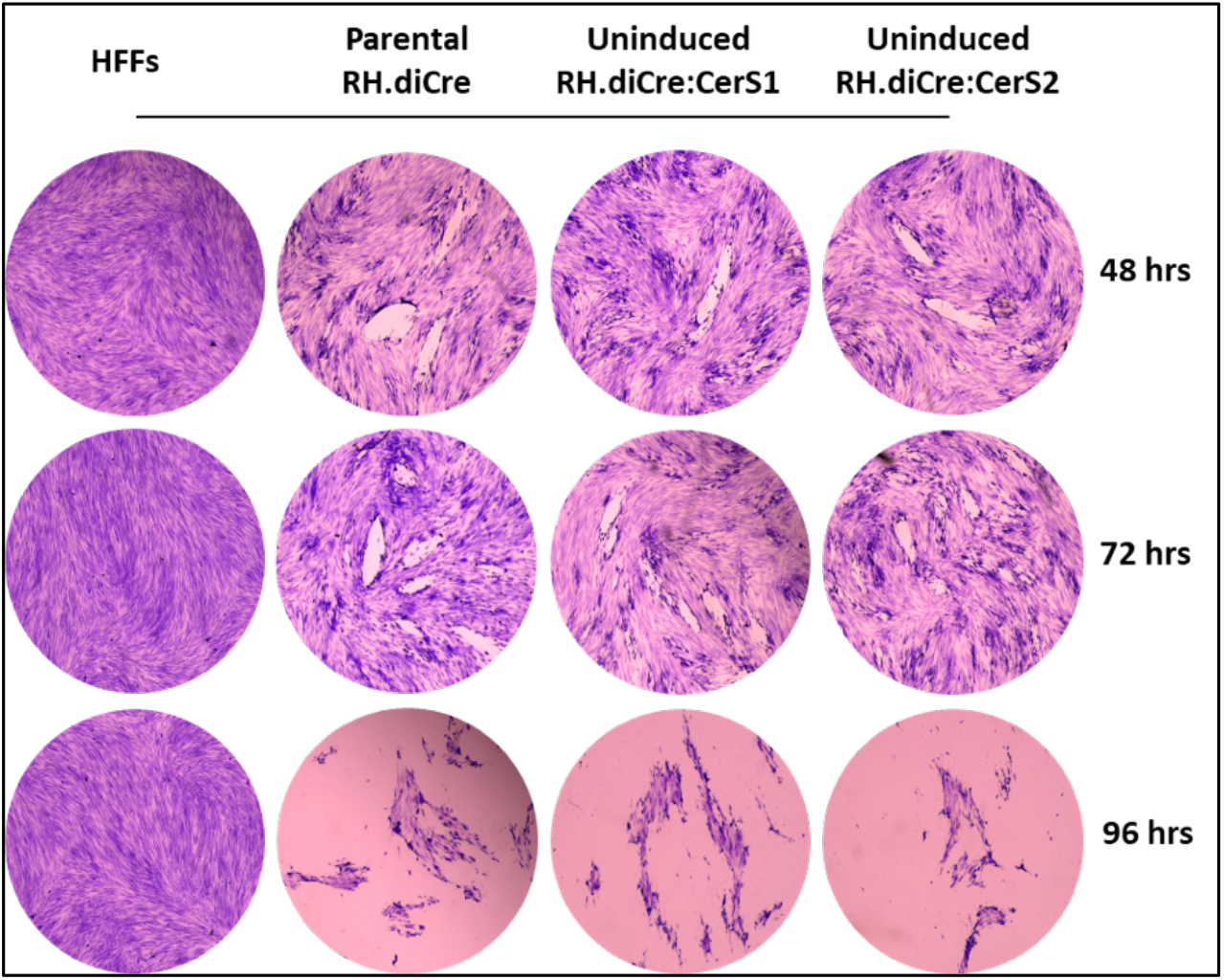
Fitness assay of parental RH.diCre (positive control), un-induced RH.diCre:CerS1 and RH.diCre:CerS2, with uninfected HFFs as a negative control. 48, 72 and 96 hours post infection.

**Fig. S5.**
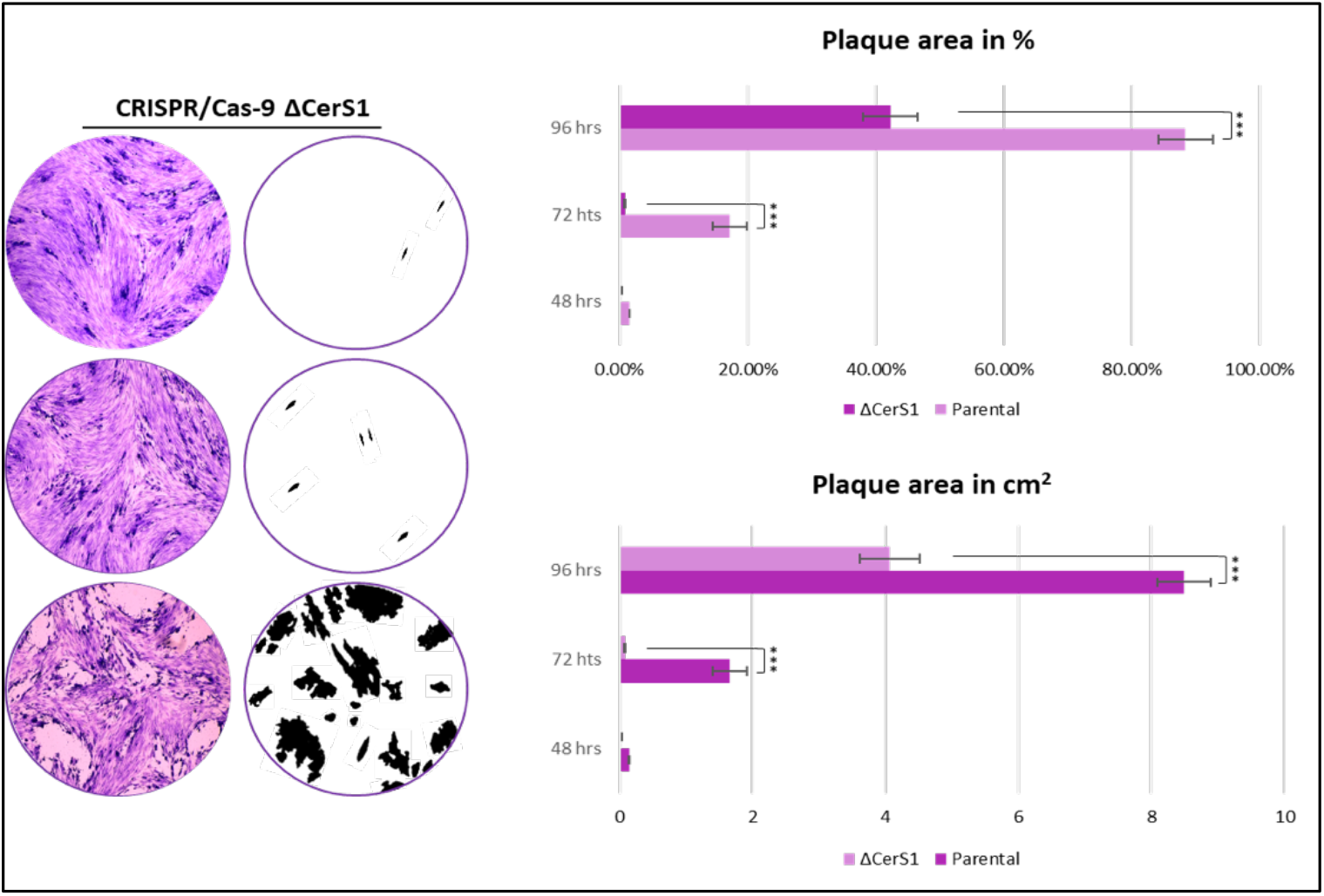
Fitness assay of CRISPR/Cas-9 *Tg*ΔCerS1 48, 72 and 96 hours post infection. Plaques are also depicted with in black for clarity. Results indicate similar parasite fitness compared to the rapamycin inducible conditional KO strain.

**Fig. S6.**
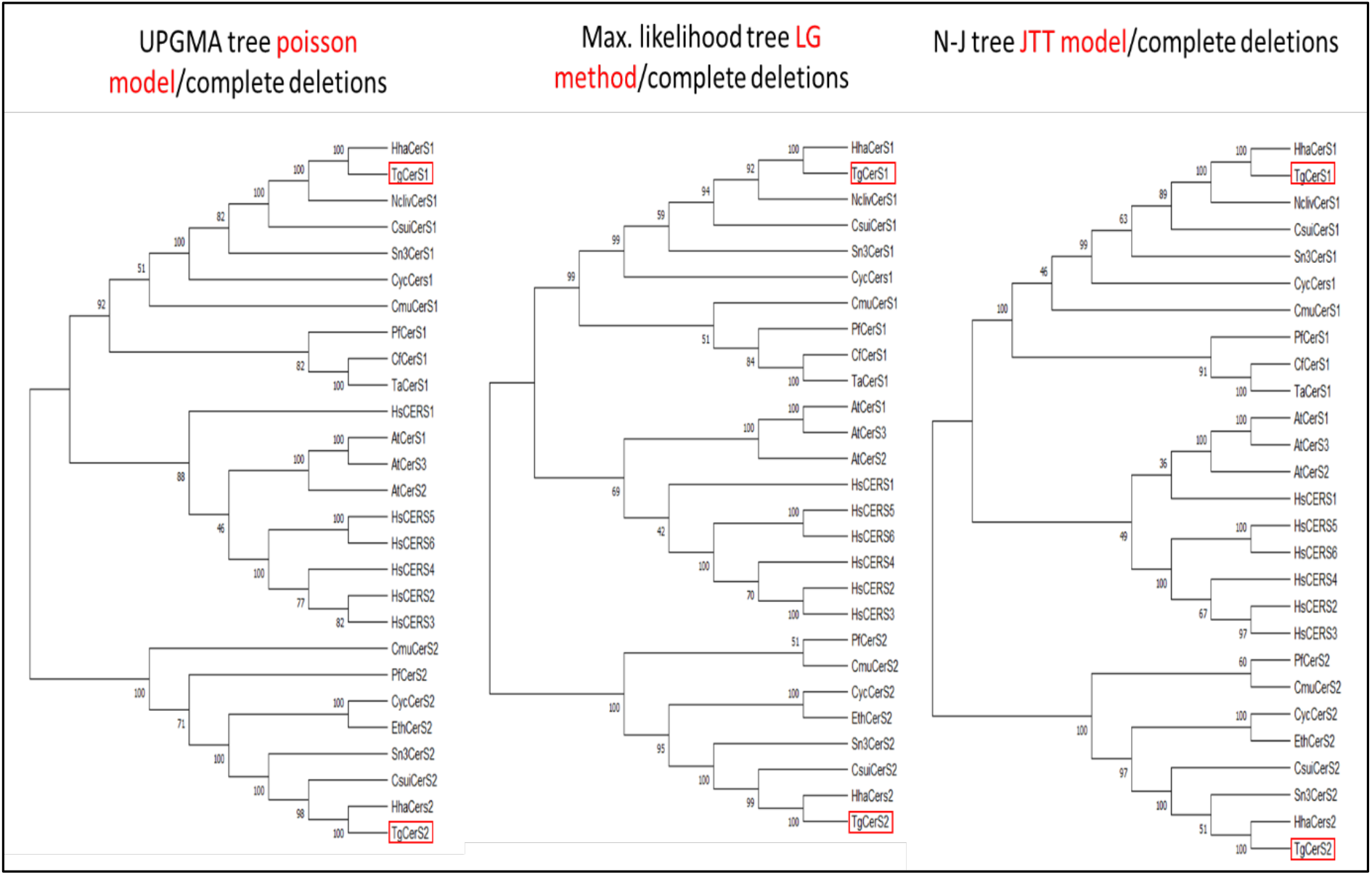
Phylogenetic trees of selected CerS orthologues as reconstructed by MEGA-X using UPGMA, Maximum likehood and N-J dedrograms based on Poisson and number of differences models. Bootstrap values are based on 1000 replicates. *Toxoplasma Tg*CerS1 and *Tg*CerS2 are highlighted in red boxes; *Tg*CerSs are highlighted in red boxes.

**Table S1.**
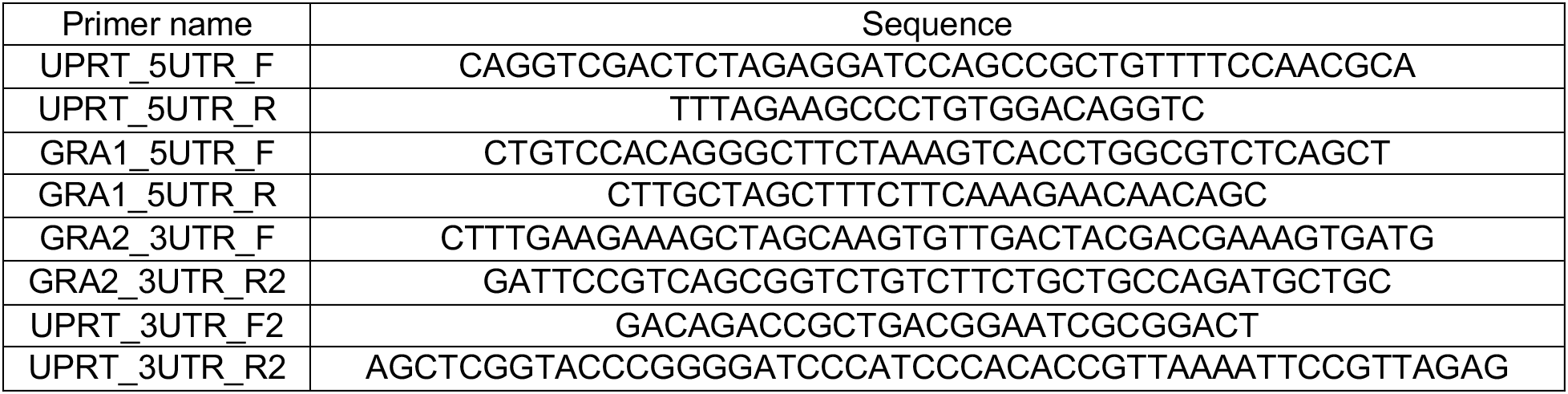
Primers used for pTOXOXpress construction. UPRT 5’ flanking region, GRA1 5’UTR, GRA2 3’UTR and UPRT 3’ flanking region were amplified from RHdeltaKu80 genomic DNA and integrated into BamHI-linearised pUC19 using in-fusing cloning.

**Table S2.**
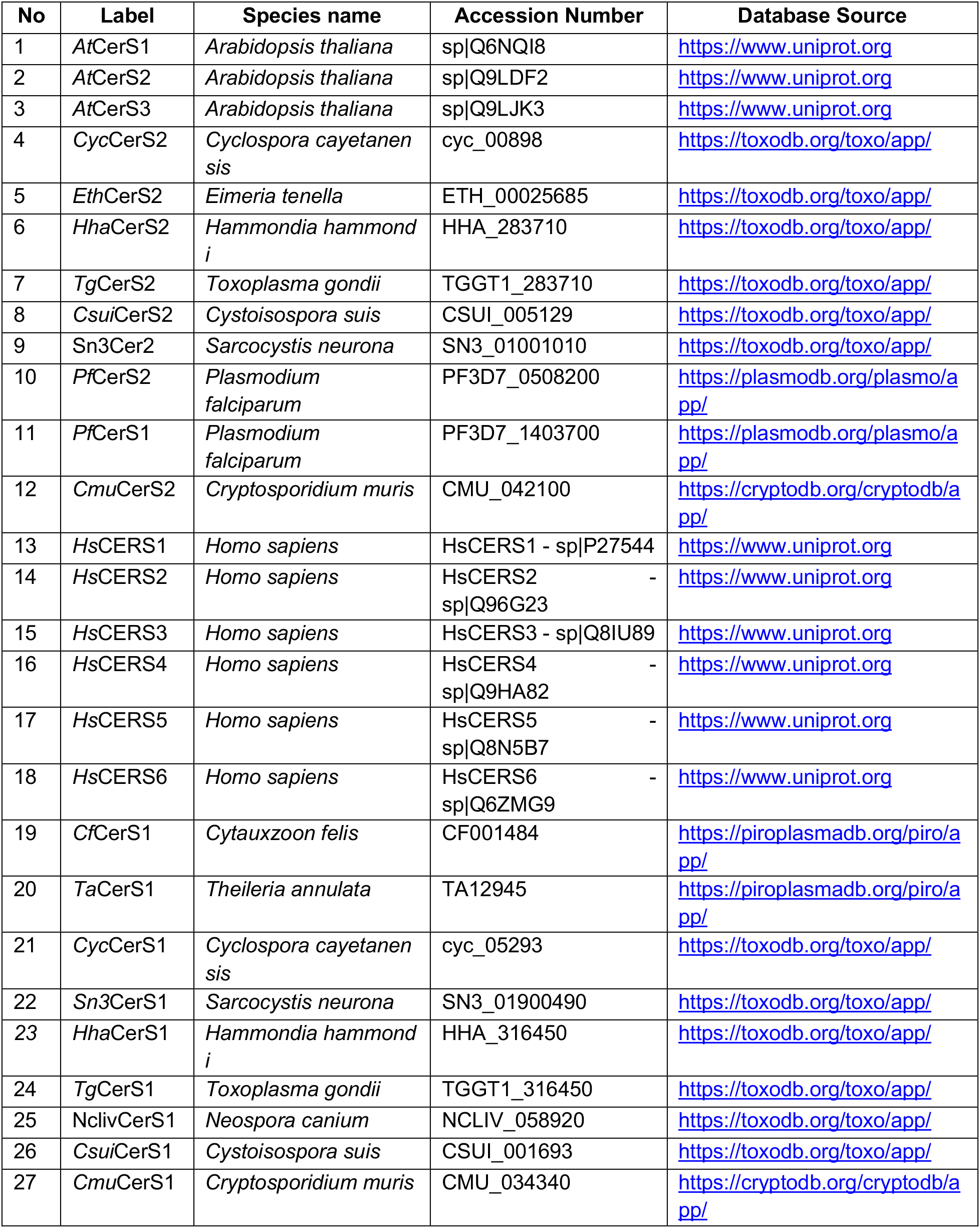
Species name, accesion numbers and online databases sources for sequences used in analyses.

**Table S3.**
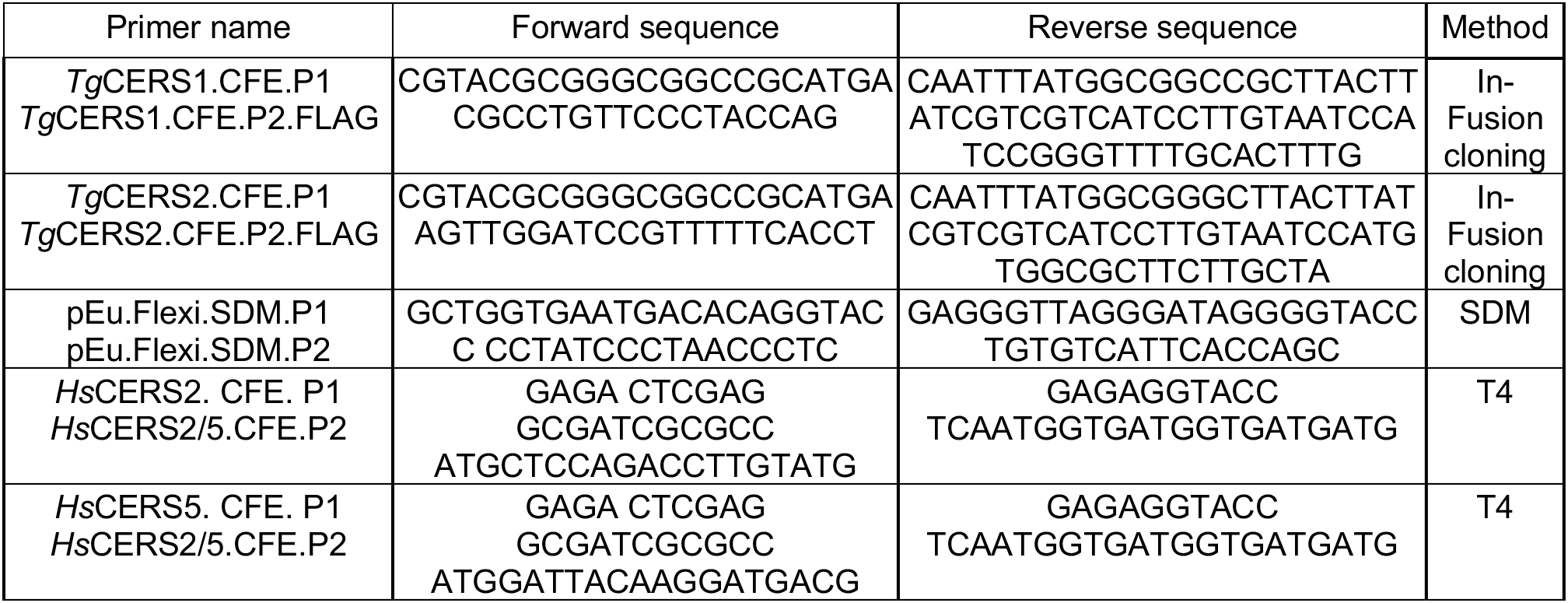
List of primers used to create plasmids for *Tg*CerS1, *Tg*CerS2, *Hs*CerS2 and *Hs*CerS5 for cell-free expression and for site directed mutatgenesis.

**Table S4.**
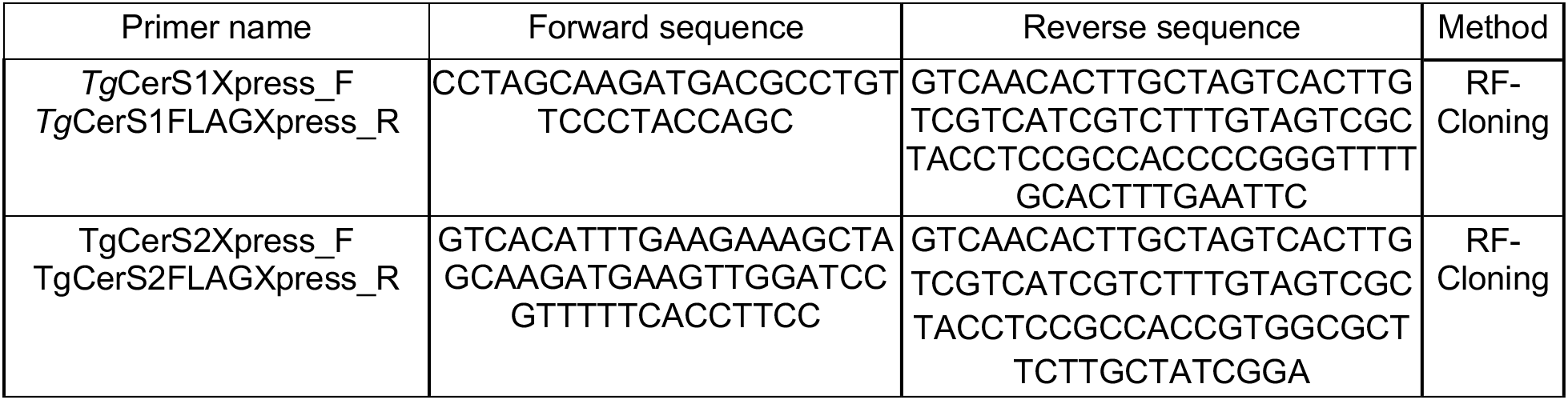
List of primers used to create *Tg*CerS1 and *Tg*CerS2 plasmids for subcellular localisation studies.

**Table S5.**
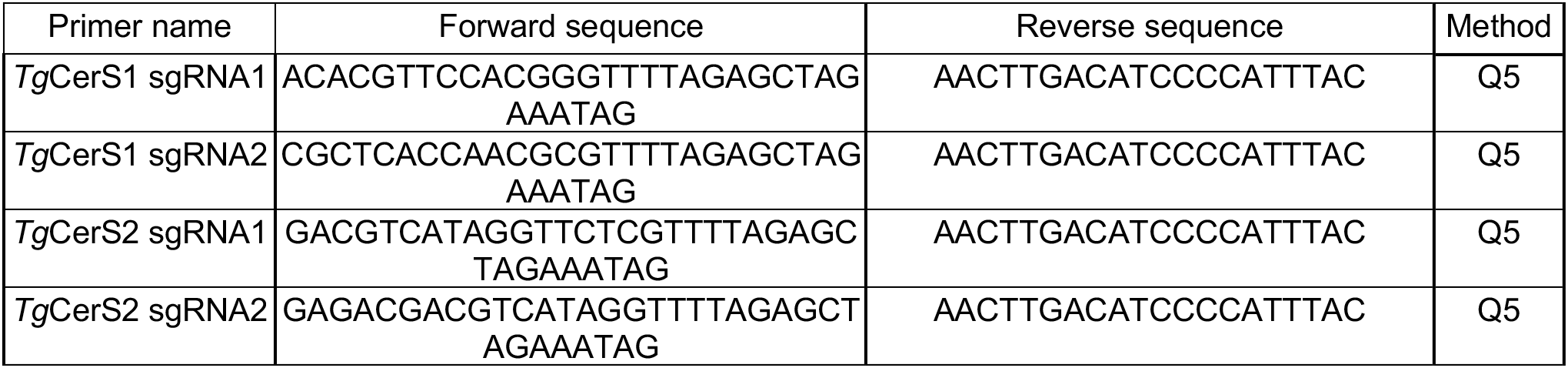
List of primers used to create CRISPR/Cas-9 plasmids. sgRNAs were desighed to target *Tg*CerS1, *Tg*CerS2 and *Tg*UPRT ORFs.

**Table S6.**
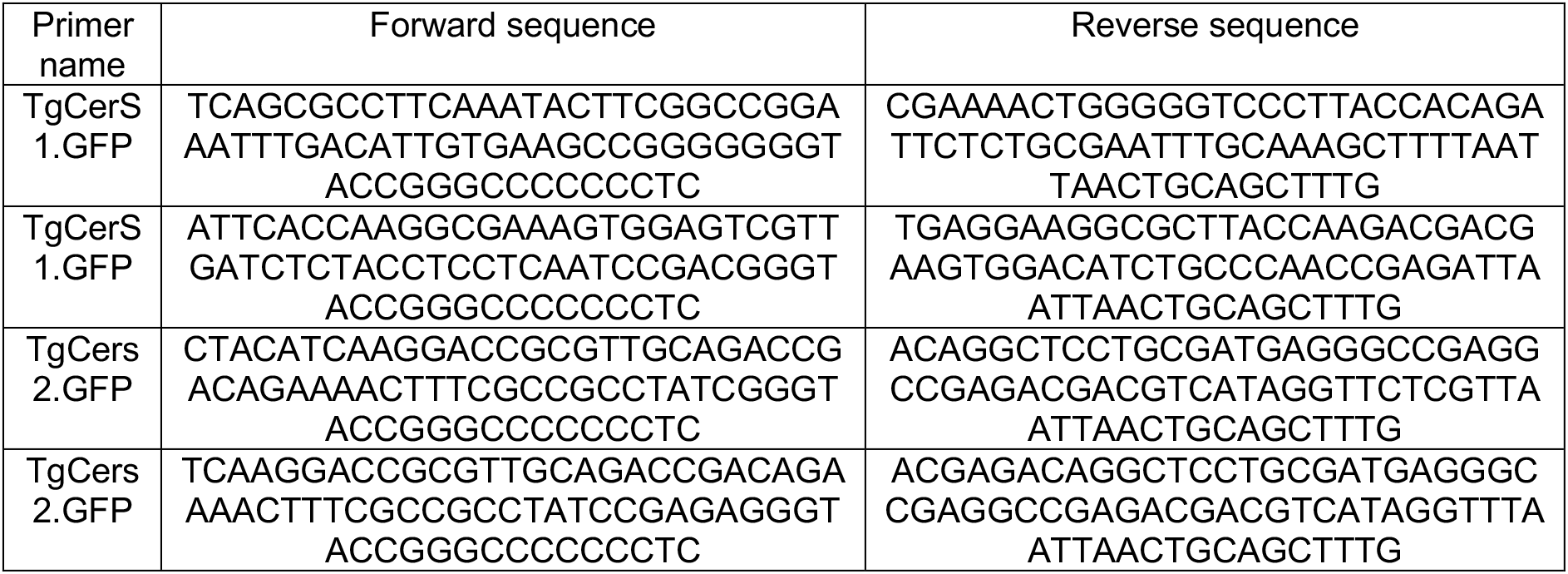
List of primers used to create GFP donor cassette. Each pair of the donor DNA primers represents 50bp flanking omology (plus overlapping region to GFP) with which the GFP cassette was amplified to undergo homologous recombination after genomic DNA was cut by CRISPR.

**Table S7.**
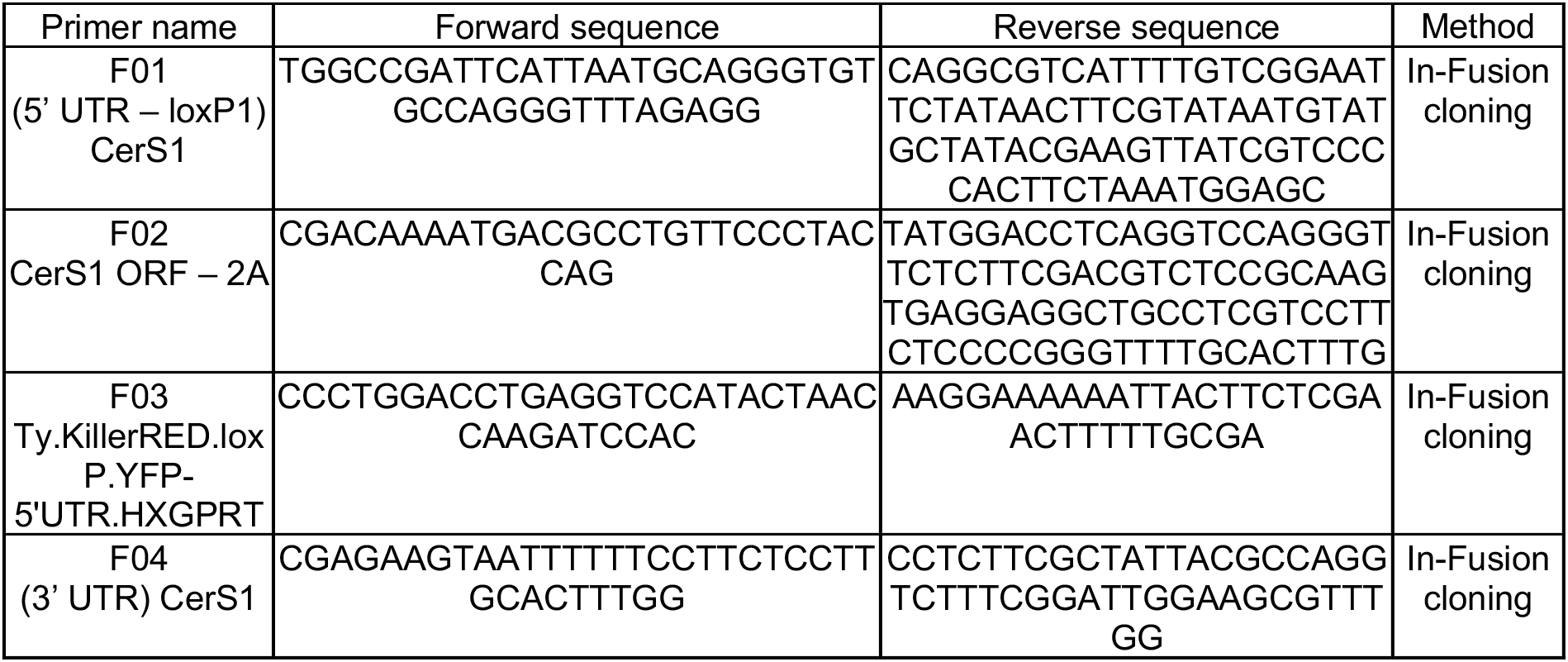
Cloning primers for diCre:*Tg*CerS1 plasmid construct.

**Table S8.**
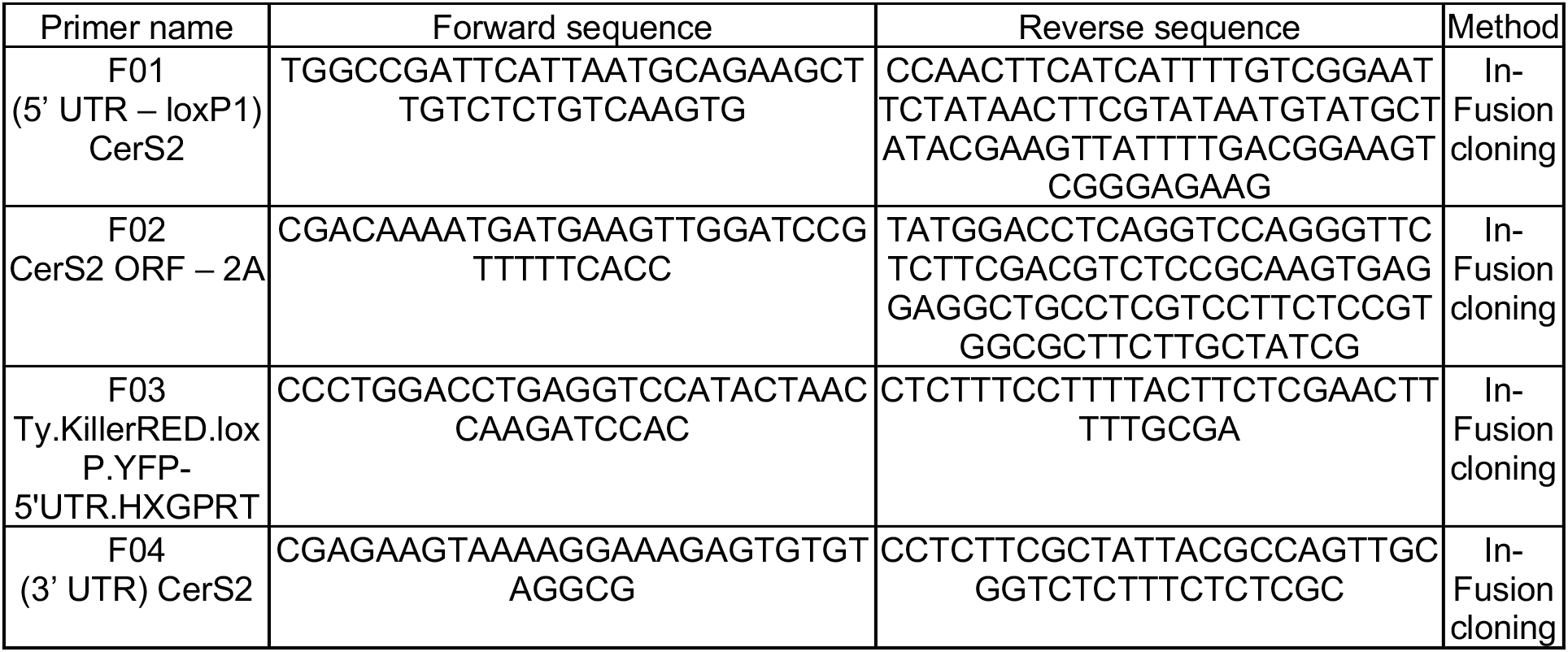
Cloning primers for diCre:*Tg*CerS2 plasmid construct.

**Table S9.**
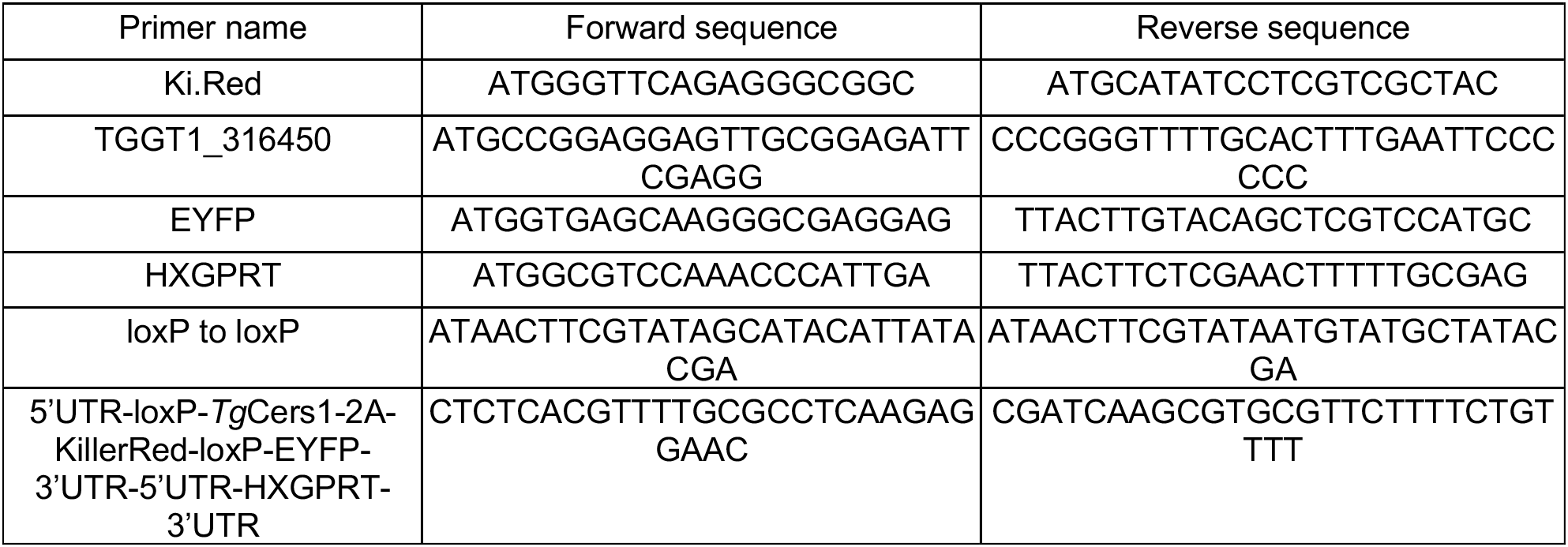
List of primers used to intentify and confirm contitional mutant strains of RH.diCre:*Tg*CerS1.

**Table S10.**
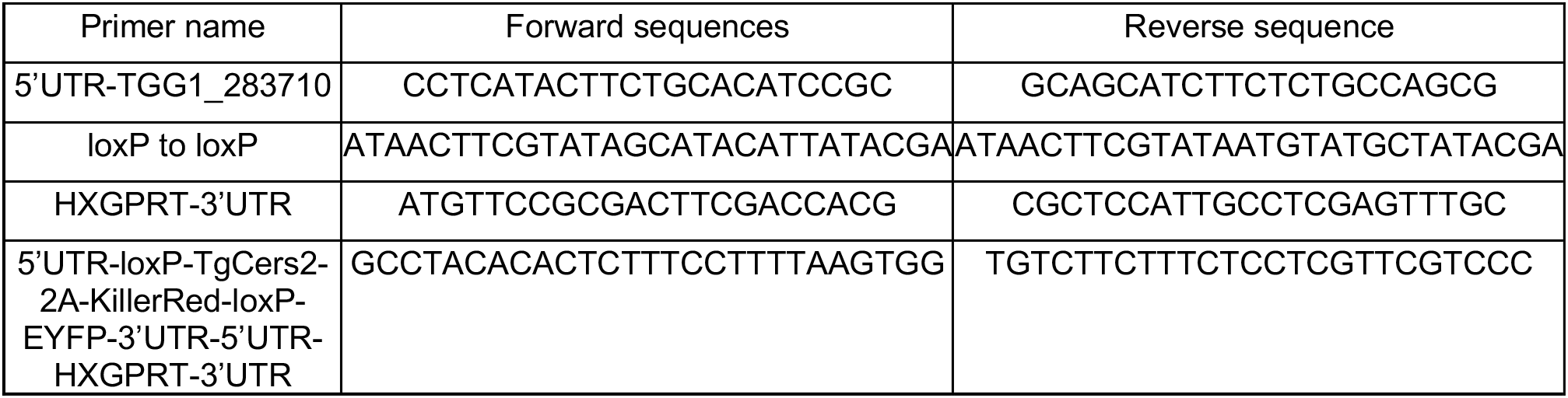
List of primers used to intentify and confirm contitional mutant strains of RH.diCre:*Tg*CerS2.

